# Hidden genetic variance contributes to increase the short-term adaptive potential of selfing populations

**DOI:** 10.1101/810515

**Authors:** Josselin Clo, Joëlle Ronfort, Diala Abu Awad

## Abstract

Standing genetic variation is considered a major contributor to the adaptive potential of species. The low heritable genetic variation observed in self-fertilising populations has led to the hypothesis that species with this mating system would be less likely to adapt. However, a non-negligible amount of cryptic genetic variation for polygenic traits, accumulated through negative linkage disequilibrium, could prove to be an important source of standing variation in self-fertilising species. To test this hypothesis we simulated populations under stabilizing selection subjected to an environmental change. We demonstrate that, when the mutation rate is high (but realistic), selfing populations are better able to store genetic variance than outcrossing populations through genetic associations, notably due to the reduced effective recombination rate associated with predominant selfing. Following an environmental shift, this diversity can be partially remobilized, which increases the additive variance and adaptive potential of predominantly (but not completely) selfing populations. In such conditions, despite initially lower observed genetic variance, selfing populations adapt as readily as outcrossing ones within a few generations. For low mutation rates, purifying selection impedes the storage of diversity through genetic associations, in which case, as previously predicted, the lower genetic variance of selfing populations results in lower adaptability compared to their outcrossing counterparts. The population size and the mutation rate are the main parameters to consider, as they are the best predictors of the amount of stored diversity in selfing populations. Our results and their impact on our knowledge of adaptation under high selfing rates are discussed.

## INTRODUCTION

Natural populations typically harbour genetic variation, especially at loci governing polygenic traits (Mittell *et al*., 2015; Wood *et al*., 2016; Clo *et al*., 2019). This variation, known as standing genetic variation, has been considered to be an important predictor for the adaptive potential of populations (Orr & Betancourt, 2001; Hermisson & Pennings, 2005; Barrett & Schluter, 2008; Pritchard *et al*., 2010; Glémin & Ronfort, 2013; Matuszewski *et al*., 2015). Indeed, standing variation represents an easily accessible source of genetic variation, that is readily available for adaptation to changing or heterogeneous conditions (Hermisson & Pennings, 2005; Barrett & Schluter, 2008). Compared to adaptation from *de novo* mutations, the probability of adapting from standing variation is higher simply because mutations already segregating in a population are expected to be present at higher frequencies (Innan & Kim, 2004; Barrett & Schluter, 2008). It has also been suggested that populations adapting from standing genetic variation can cope with more severe and more rapid environmental change, as they are able to cross larger distances in the phenotype space (Matuszewski *et al*., 2015). The amount of standing variation available in a population is thus expected to play a key role in adaptation, and any forces affecting it may greatly influence whether or not populations are able to survive new environments.

The mating system is a population characteristic that is known to greatly affect the amount and structure of genetic variation. For instance, both theoretical models (Charlesworth & Charlesworth, 1995; Lande & Porcher, 2015; Abu Awad & Roze, 2018) and empirical data (Charlesworth & Charlesworth, 1995; Geber & Griffen, 2003; Clo *et al*., 2019) have shown that, compared to outcrossing populations, self-fertilization reduces, on average, the amount of additive genetic variance for polygenic or quantitative traits under stabilizing selection. This diminution is due to more efficient purifying selection under selfing and to negative linkage disequilibria maintained between alleles at different loci: the so-called Bulmer effect (Lande & Porcher, 2015; Abu Awad & Roze, 2018). Due to this reduction in genetic variability, predominant selfing has been described as an evolutionary dead-end (Stebbins, 1957; Takebayashi & Morrell, 2001; Igic & Busch, 2013). However, recent theoretical work and empirical data have highlighted that cryptic genetic variability could contribute to the adaptive potential of natural populations. Cryptic genetic variation has been defined as the part of a population’s standing genetic variation that has no effect on the phenotypic variation in a stable environment but can increase heritable variation in environmental conditions rarely experienced (Gibson & Dworkin, 2004; Paaby & Rockman, 2014). Such variability has been detected in both outcrossing (in sticklebacks, McGuigan *et al*., 2011, cavefish, Rohner *et al*., 2013, dung flies, Berger *et al*., 2011, gulls, Kim *et al*., 2013 or spadefoot toads, Ledon-Rettig *et al*., 2010) and selfing species (*Caenorhabditis elegans*, Milloz *et al*. 2008; *Arabidopsis thaliana*, Queitsch *et al*. 2002). Two main mechanisms could explain the accumulation and the release of such variance: interactions between loci (Badano & Katsanis, 2002; Shao *et al*., 2008), and phenotypic plasticity (Anderson *et al*., 2013). In this paper, we focus on interactions between loci maintained at stabilizing selection.

In maintaining the population close to the phenotypic optimum, stabilizing selection disfavors genetic and phenotypic diversity (Lande & Porcher, 2015; Abu Awad & Roze, 2018). However, the structure of the additive variance also strongly depends on the trait mutation rate and the prevalence of pleiotropy (Charlesworth, 1990; Lande & Porcher, 2015; Abu Awad & Roze, 2018). When the per-trait mutation rate is weak, associations between loci are negligible (mutations of strong effect arise slowly and are easily purged), but when the rate increases, the creation and maintenance of co-adapted gene complexes structure the additive variance into positive within-loci components and negative among-loci components of variance, reducing the observed level of additive variance (Abu Awad & Roze, 2018). The remobilization through recombination of this among-loci component of variance could boost the adaptability of populations undergoing an environmental shift (Le Rouzic and Carlborg 2008). Indeed, if associations between loci are broken, segregating alleles could express some or all of their additive effects in new genetic backgrounds. Such remobilization is only possible if residual allogamy occurs (*i*.*e*. if the selfing rate is not equal to 1), which is expected to be common in natural plant populations (Kamran-Disfani & Agrawal, 2014; Clo *et al*., 2019). Classical models analyzing the effect of selfing on adaptation from standing genetic variation have considered a single locus (Glémin & Ronfort, 2013), thus neglecting interactions among loci that could contribute to an important fraction of standing genetic variation. As self-fertilization reduces the effective recombination rate (Nordborg, 2000), allowing the maintenance of co-adapted gene complexes, the storage of genetic diversity through genetic associations should be more prevalent in selfing populations (as suggested in Charlesworth, 1990; Lande & Porcher, 2015; Abu Awad & Roze, 2018).

In this paper, we explore this hypothesis, using a quantitative genetics framework. We describe and quantify how, to what degree, and under which conditions populations accumulate genetic variation at polygenic traits. Because the development of tractable analytical predictions of the phenotype and genetic variance dynamics is not trivial and often requires strong hypotheses to be made, notably random mating and considering the system to be at linkage equilibrium (Barton & Turelli, 1987; Keightley & Hill, 1989; Bürger, 1993), we chose to use a simulation approach. Although it is known that directional dominance plays an important role in the fate of mutations, especially in the presence of self-fertilization, the interaction of both directional dominance and epistasis with mating systems are not well known. In order to avoid further complicating the interpretation of our results, we have considered a fully additive model, which despite its simplicity, has previously led to surprisingly accurate predictions (Martin *et al*., 2007; Manna *et al*., 2011). We show that, in models allowing genetic diversity to be stored through genetic associations and when adaptation is only possible from pre-existing standing genetic variation, predominantly selfing populations can adapt as well as their mixed-mating and outcrossing counterparts, when mutation rate is high (but realistic), despite initially low levels of observable genetic diversity. If the mutation rate is low, purifying selection impedes the storage of diversity through genetic associations, and our simulations confirm the more classical results that selfing populations adapt less well than their outcrossing counterparts do.

## MATERIAL AND METHODS

### General assumptions

We consider the evolution of a quantitative trait *Z* in a population of size *N*, made of diploid individuals reproducing through partial self-fertilization, with a constant selfing rate, (and hence constant inbreeding level) *σ*. The phenotypic value *z* of an individual is determined by the additive action of *L* loci each with an infinite possible number of alleles and is given by

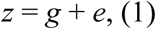

where *g* is the genetic component of the individual’s phenotype, and is given by 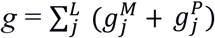, with *g*^M^_*j*_ (respectively *g*^P^_*j*_) the additive allelic effect at locus *j* inherited from the maternal (respectively paternal) gamete. There are no dominance or epistasis at the phenotypic scale in this model, but both of which arise naturally at fitness scale, when considering stabilizing selection and a mean phenotype close to the optimum, the mean dominance h ≈ 0.25 and epistasis effects are on average null for fitness. The random environmental effect, *e*, is drawn from a Gaussian distribution of mean 0 and variance *V*_E_, and is considered to be independent from the genetic components of fitness. The trait initially undergoes stabilizing selection around an optimal phenotypic value, denoted *Z*_opt_. The fitness value *W*_Z_ of an individual with phenotype *z* is thus described by the Gaussian function:

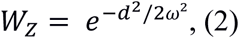

where *d* is the distance between the individual’s phenotype *z* and the optimum trait value and *ω*^2^ is the width of the fitness function, and represents the strength of selection.

### Simulation model

We implement the model described above into an individual based simulation model written in C++, a modified version of the “continuum of alleles” program provided in Abu Awad and Roze (2018) available in supplementary materials and online (https://github.com/dialaAbAw/SelfingAdaptation).

We consider a population of *N* diploid individuals, each represented by two linear chromosomes with *L* multi-allelic loci, coding for a single quantitative trait under selection. At the beginning of each simulation, all individuals are genetically identical and are at the phenotypic optimum (all loci carry alleles with effect 0 and *Z* _OPT_ = 0). The life cycle can be summarized by five successive events. First, there is a phenotype-dependent choice of the first parent (selection), followed by mating-type choice (selfing versus outcrossing at rates σ and (1-σ) respectively) and then, in the case of outcrossing, phenotype-dependent choice of the second parent. Selection takes place as follows: if the ratio of the selected parent’s fitness over the highest recorded fitness value in the current generation is higher than a number sampled in a uniform law comprised between 0 and 1, the individual is allowed to reproduce. Once the two parents are chosen, they each contribute a gamete, produced through uniformly recombining the parental chromosomes. The number of cross-overs is sampled from a Poisson distribution with parameter *R*, the map length. From Haldane’s mapping function, the recombination rate between two adjacent loci is 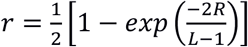. We choose parameters that ensure that *r*≈ 0.5, such that loci remain, at least physically, unlinked. This phase is then followed by the introduction of new mutations, the number of which is sampled from a Poisson distribution with parameter *U* (with *U* = *µL, µ* being the per locus mutation rate). The additive value of a new mutant allele is drawn from a Normal distribution of mean 0 and variance *a*^2^.

Each simulation consists of two phases, the first being burn-in time to allow the population to attain Mutation-Selection-Drift equilibrium (M-S-D). The population is considered to be at M-S-D equilibrium when the average fitness value calculated over the last thousand generations does not differ by more than one percent from the mean fitness calculated over the previous thousand generations. The second phase consists of following the population after a shift in the phenotypic optimum. After the shift, the haploid genomic mutation rate *U* is set to 0 so that the only source of genetic variability to reach the new optimum is the standing variation accumulated before the environmental change.

### Simulation parameter values

Simulations were run for several parameter sets in order to evaluate the effects of each parameter on both the equilibrium conditions and on the ability of populations to adapt after the environmental change. The values chosen for the mutation rate *U* range from 0.005 to 0.1, reflecting the *per*-trait haploid genomic mutation rate found in the literature (Keightley & Bataillon, 2000; Shaw *et al*., 2002; Haag-Liautard *et al*., 2007). We use parameter set values similar to those explored in Bürger *et al*., (1989) and Ronce *et al*., (2009), with the number of freely recombining loci under selection *L* = 50, and *a*^2^ = 0.05, *V*_E_ =1, *ω*^2^ = 1, 9 or 99. In this model, the mean deleterious effect of mutations 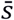 is therefore equal to 0.0125, 0.0025 or 0.00025 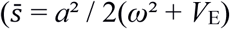, Martin and Lenormand 2006).

Before the shift in the optimum, *Z* _OPT_ is set to 0, and after M-S-D equilibrium it is set to 2.5, so that the shift is of order *L*.*a*^2^, and ensure that adaptation is due to genetic changes and not environmental effects (Ronce *et al*., 2009). We follow populations only over 20 generations after the shift in the optimum, as we chose to follow the possibility of adaptation without any contribution from *de novo* mutations. After this time limit, genetic variation starts to erode, and the observed results are mainly due to the effects of loss of diversity due to drift and not to the selection process. Although simulations were run over a large range of selfing rate values, throughout the manuscript we show results run principally for three rates of self-fertilization, *σ* = 0, 0.5 and 0.95, representing outcrossing, mixed-mating and predominantly selfing respectively. These three values were chosen because they were representative of the main patterns of adaptation we observed over the whole range of selfing rates (*σ* from 0 to 1). We also considered three population sizes *N* = 250, 1000 or 10000, which allow to test the effect of drift.

#### Analysis of the effect of selfing on the equilibrium genetic variance

Following Turelli & Barton (1990), we decompose the genetic variance of a polygenic trait using the following equation:

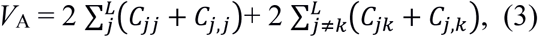

with

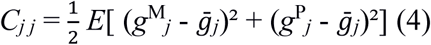

and

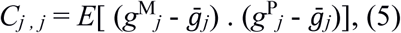

where 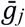 is the mean allelic effect on the phenotype at locus *j* and *g*^M^_*j*_ (respectively *g*^P^) is the allelic effect at locus *j* inherited from the maternal (respectively paternal) gamete. The sum of of *C*_*jj*_ over loci represents the genic variance (noted *V*_genic_, which is the genetic variance of a trait in a population harboring the same allelic frequencies as the population under study, but without any genetic associations between loci), and is computed from simulation outputs following equation (4). The sum of all values of *C*_*j,j*_ represents the covariance in allelic effects at locus *j* on the maternally and paternally inherited chromosomes, and represents the fraction of the genetic variance due to excess of homozygosity (noted *V*_inbred_); we compute it following equation (5). This quantity represents *F*.*V*_genic_, where *F* is the inbreeding coefficient of the population. The first term of equation (3) 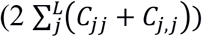 represents the genetic variance due to within locus variation. The second term 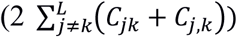 represents the component of the variance due to associations between loci (noted *COV*_LD_), and is obtained by computing the sum of covariances between genotypic values among all pair of loci. This component is proportional to linkage disequilibrium (LD), and tends to be negative under stabilizing selection due to associations between alleles at different loci with compensatory effects (*i*.*e*. the allele on one locus is positive, the other negative, their effects on the phenotype thus cancel out when both are present). At any time in the simulated populations, *V*_A_ = *V*_genic_ + *V*_inbred_ + *COV*_LD_.

As a check, estimates of additive genetic variance observed at M-S-D equilibrium in our simulations were compared to analytical predictions obtained by Bürger et al. (1989) (Stochastic House of Cards approximate) and by Abu Awad and Roze (2018) under similar assumptions. Comparisons and details are shown in Appendix A.

#### Analyses of the response to an environmental change

To analyse the population’s response to an environmental change, we focus on the 20 first generations after the environmental change. We follow the temporal dynamics of the additive genetic variance of the trait and of its components. We also analyze changes with time of the mean trait value and population fitness, as well as the dynamics of the number of haplotypes and the proportion of new haplotypes generated after the environmental change.

In addition, and in order to determine if the remobilization of *COV*_LD_ plays a role in the adaptive process of selfing populations, we compute for each mutation rate, the slope of the trait mean variation shortly after the environmental change (during the first five generations) as a function of the amount of additive variance available at M-S-D equilibrium. If remobilization of *COV*_LD_ is involved in the adaptive process of selfing populations, the initial slope, for an initially similar amount of additive variance, should be higher in selfing populations than in mixed mating and outcrossing ones.

## RESULTS

### LEVEL AND STRUCTURE OF ADDITIVE GENETIC VARIANCE AT MUTATION SELECTION DRIFT EQUILIBRIUM

In agreement with expectations from previous works, the additive genetic variance present at M-S-D equilibrium in our simulations is overall negatively correlated with the selfing rate (Figure 1), in part due to the higher efficiency of selfing in purging deleterious mutations. In general, reducing the strength of stabilizing selection increases the overall amount of additive variance, as mutations of small effect are not as easily purged from the population (Figure 1). The population size, however, has little qualitative effect on the relationship between selfing and *V*_A_, at least as long as the mutation rate is not too high (Figure S1). As detailed in Appendix A, the values of *V*_A_ obtained from our simulations were compared to expectations from analytical approximations from Abu Awad & Roze (2018) and the Stochastic House of Cards approximation (SHC, Bürger *et al*., 1989). We generally find a good agreement between our results and the SHC approximation for the smaller population sizes and outcrossing populations (*σ* = 0 or 0.5). Approximations from Abu Awad & Roze (2018) only hold for predominantly selfing populations, considering the whole genome as a single locus (*σ* = 0.95, equation 44 in Abu Awad & Roze, 2018), or for all mating systems when population size is large (*N*=10.000) and the strength of selection weak (*ω*^*2*^=99, equation D23). The lack of fit between our results and approximations from Abu Awad & Roze (2018) is probably due to the small number of loci considered in our simulations (*n* = 50 in our parameter set) and to the fact that we consider only a single trait, both parameters violate the assumptions for these analytical approximations to hold.

**Figure 1.**
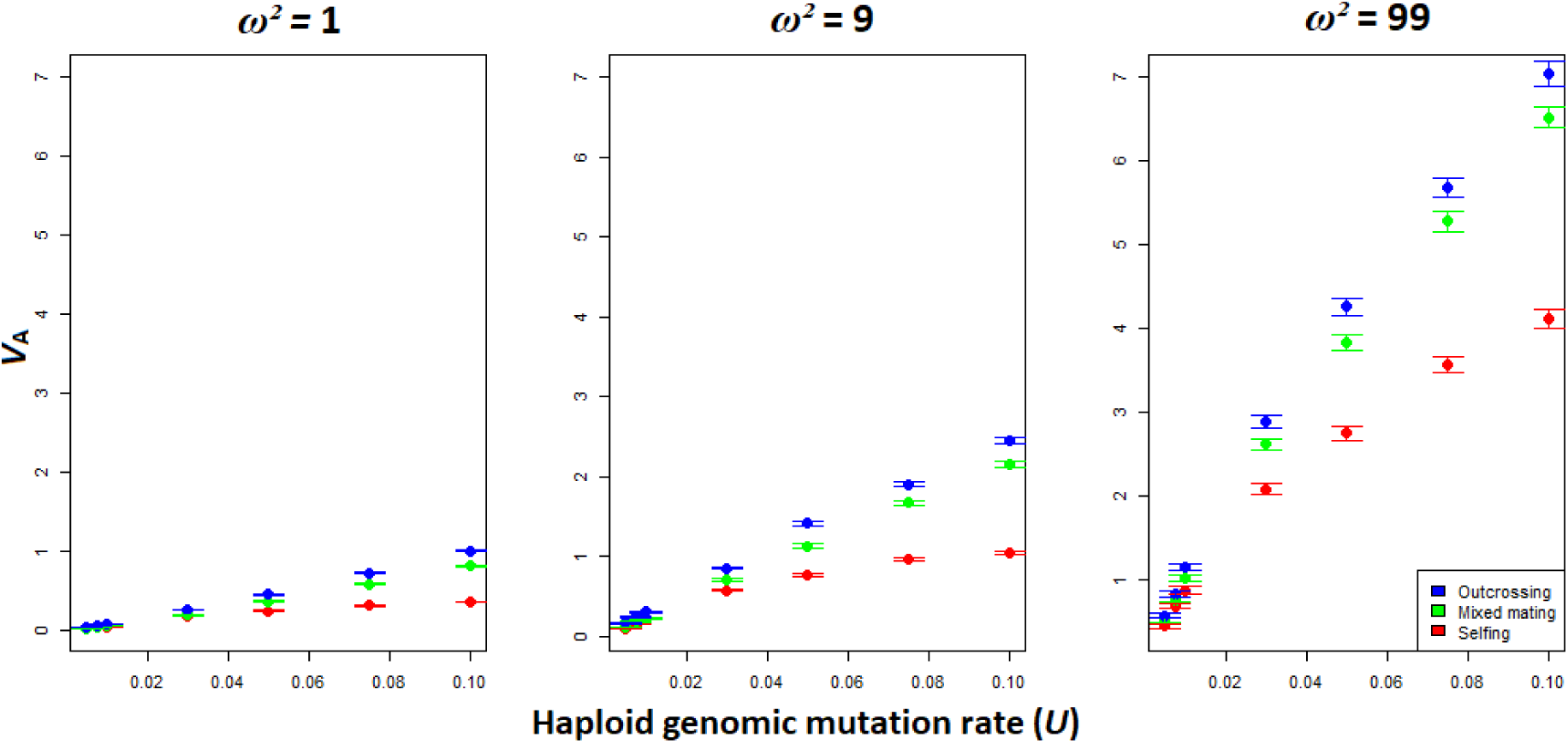
Additive genetic variance observed at mutation-selection-drift equilibrium in the simulated populations, for different mating systems, *N*=1000, and different strengths of selection (from *ω*^*2*^=1 (strong selection) to *ω*^*2*^=100 (weak selection), from the left to the right). Error bars stand for the 95% confidence interval (*n* = 100 simulations).

By decomposing the observed additive variance *V*_A_ into the components described in the Methods above (*V*_genic_, *V*_inbred_, *COV*_LD_), we find that for the range of parameters we chose to explore, the strong decrease in *V*_A_ observed under predominant selfing is mostly driven by a large but negative *COV*_LD_ (Figure 2). This confirms the expectation that recurrent selfing favours the maintenance of associations among loci. Under such associations the efficiency of purifying selection is reduced, as the selection pressure on a single mutation is weakened by its positive association with other mutations at other loci, which in turn contributes to higher *V*_genic_ and *V*_inbred_ in selfers compared to outcrossers, as observed in Figure 2. Our analysis also showed that the higher the mutation rate, the more genetic associations are maintained and the weaker the efficiency of purifying selection. The increase of both *V*_inbred_ and *V*_genic_ can, in some cases, overcome the negative effect of *COV*_LD_, as is the case for large populations (*N*=10.000), leading to selfing populations that have close to, and sometimes even greater amounts of genetic variance than outcrossing or mixed mating populations (Figure S2). As can be expected from classical population genetics models, smaller population size leads to a decrease in V_A_, whereas higher mutation rates tend to increase it (Figure S1).

**Figure 2.**
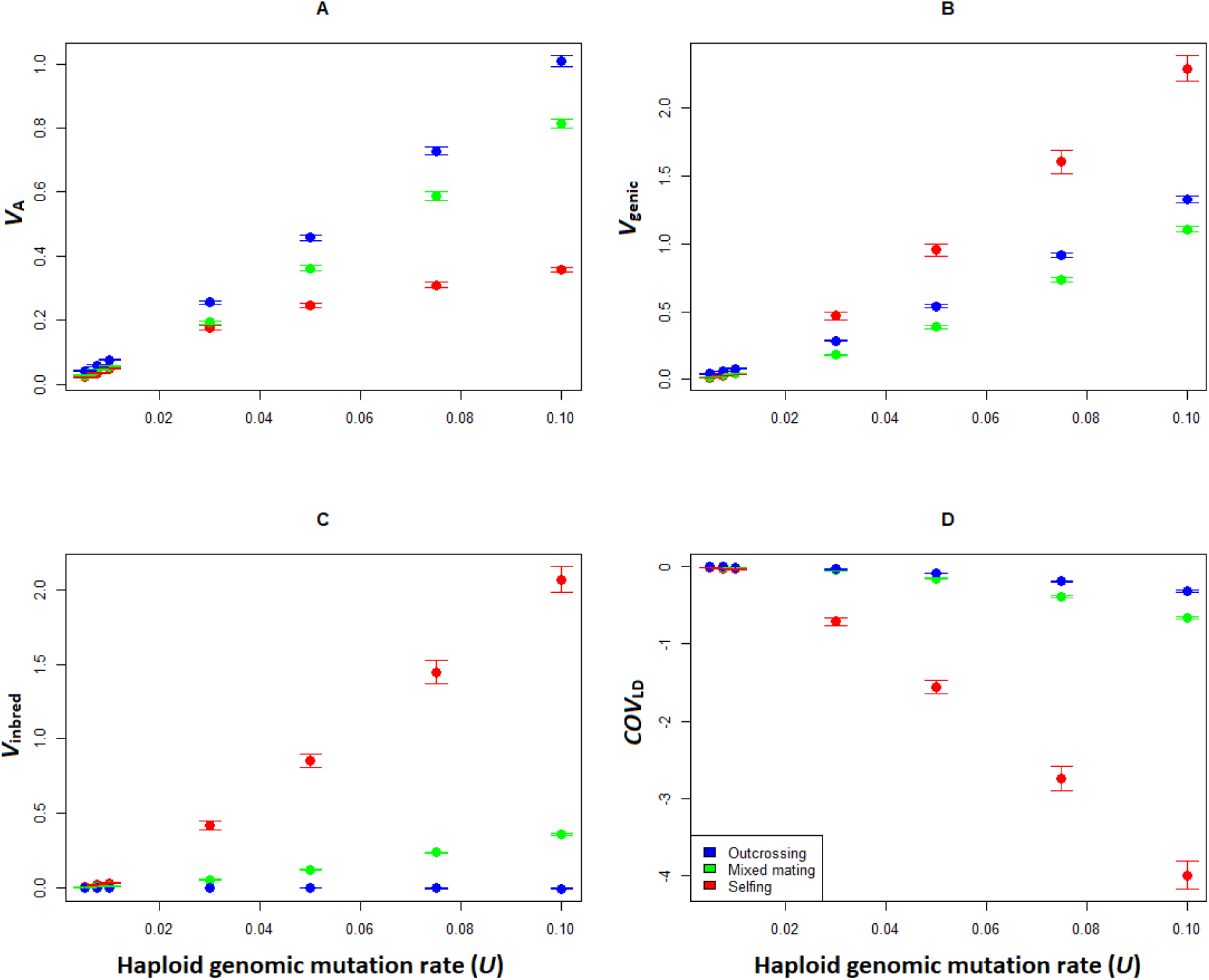
Total additive genetic variance and its three components estimated at MSD equilibrium, for different genomic mutation rates and mating systems, for *N*=1000 and *ω*^*2*^=1. **A**. Total additive variance of the phenotypic trait, identical to Figure 1 first panel. **B**. Genic variance for the phenotypic trait (*V*_genic_). **C**. Genetic variance due to inbreeding (*V*_inbred_). **D**. Genetic covariance due to linkage disequilibrium (*COV*_LD_). Error bars stand for 95% confidence interval (*n* = 100).

### PATTERNS OF ADAPTATION THROUGH STANDING GENETIC VARIATION

After populations have reached mutation-selection-drift balance, an environmental change is induced and no new mutations are introduced. Population dynamics are followed over 20 generations and, if during this time the population reaches a similar level of fitness as that observed before the environmental change, then we consider that the population is able to adapt to the new optimum. When the strength of purifying selection is moderate to weak (*ω*^*2*^ = 9 or 99), all populations seem to harbour enough genetic variation to quickly and efficiently respond to an environmental change, independently of their selfing rate. As there were no observable dynamic changes in *V*_A_ and its components for these parameter values over the course of the first 20 generations following the environmental change, the results for these simulations are only presented in Supplementary Figures S3 to S8. In the following sections we will only be concentrating on the case of *ω*^*2*^ = 1, representing the strongest selection pressure we explored for.

### DYNAMICS OF PHENOTYPE AND FITNESS DURING ADAPTATION UNDER STRONG SELECTION

As illustrated in Figure 3, when the mutation rate is small (*U* = 0.005), and hence the amount of diversity stored through genetic associations for selfing population is also small, none of the populations are able to fully adapt during the 20 first generations, and population fitness remains low. In this case the amount of within-loci genetic variation at equilibrium is the only source of diversity, and as outcrossing and mixed mating populations have higher *V*_A_, they reach phenotypic values closer to the new optimum, and have a significantly higher fitness than their selfing counterparts (Figure 3 for *U*=0.005). For higher mutation rates (*U* = 0.1), dynamics of the phenotypic trait and fitness are extremely similar for all mating systems, despite lower genetic diversity in selfing populations. As shown in the following sections, when the mutation rate increases, the variance stored in selfing populations through genetic associations is partially released, increasing the adaptive potential of these populations. In such cases, selfing populations are able to reach (1) the new phenotypic optimum, and (2) level of fitness similar to those observed at M-S-D equilibrium, thus performing as well as outcrossing populations (Figure 3).

**Figure 3.**
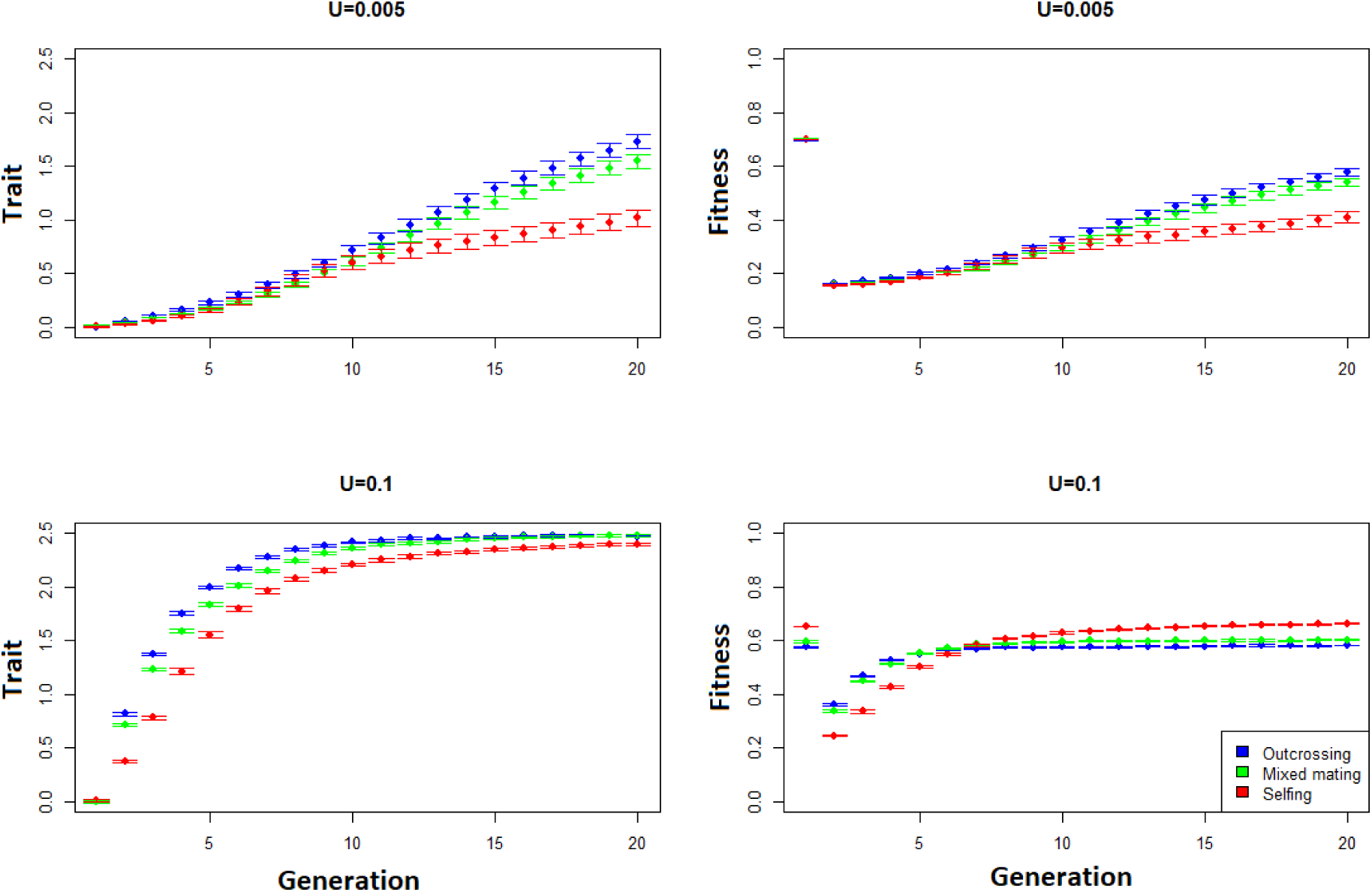
Dynamics of the mean trait value and of the mean fitness value of the population during the 20 generations following the environmental change, for different haploid mutation rates (Top-down: *U*=0.005; *U*=0.1) and as a function of the mating systems. Other parameter values are *N*=1000 and *ω*^*2*^=1. Error bars stand for 95% confidence interval (*n*=100).

The above-mentioned patterns are similar irrespective of the population size. As mentioned in the first section of the results, the greater the population size the greater the genetic diversity (*V*_A_ and *COV*_LD_ for selfing species) for a given mutation rate, increasing the range of mutation rates for which selfing populations are at least as well, if not better, adapted as outcrossing and mixed mating populations (Figures S9 and S10). This means that population size has a quantitative but not a qualitative effect.

### DYNAMICS OF THE ADDITIVE VARIANCE AND OF ITS COMPONENTS

As the population undergoes directional selection, the dynamics of the observable additive variance and each of its components are telling of the processes driving adaptation to the new optimum. In small outcrossing populations (*N* = 250), the additive variance exhibits very small changes during the adaptation process, as do its components, whatever the strength of selection (Figures S9 to S11). For larger population sizes (*N*=1000 or 10.000), the additive variance increases during the first generations when the mutation rate is small to medium (*U* ≤ 0.05, Figure 4A for *N*=1000, see supplementary materials for *N*=10.000). This increase of *V*_A_ stems from an increase in *V*_*genic*_, indicative of the change in the phenotype being driven by an increased frequency of alleles that may have been neutral or deleterious before the environmental change (Bürger 1993). For high mutation rates (*U* = 0.1), as for small populations, the additive genetic variance and its components remain constant through time (Figure 4). *V*_inbred_ does not vary (Figure 4C). Overall patterns are similar for mixed mating populations.

**Figure 4.**
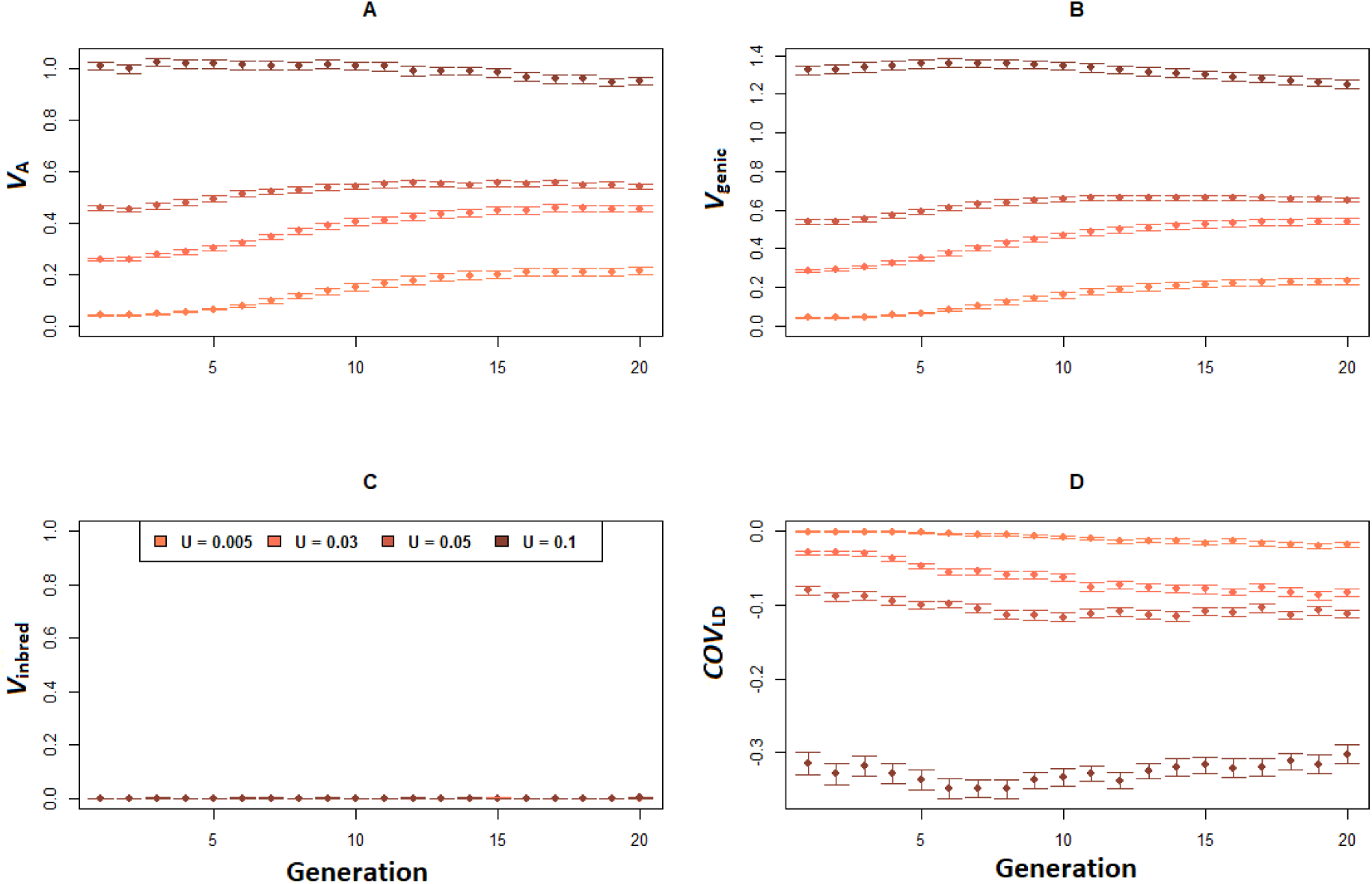
Dynamics of the additive genetic variance and its components in outcrossing populations (*σ*=0), mutation rates are set at zero after generation 0. **A**. Observed additive variance for the phenotypic trait. **B**. Genic variance for the phenotypic trait (*V*_genic_). **C**. Genetic variance due to inbreeding (*V*_inbred_). **D**. Genetic covariance due to linkage disequilibrium (*COV*_LD_), for different mutation rates (U=0.005; U=0.03; U=0.05; U=0.1). Error bars stand for 95% confidence interval (n = 100). Other parameter values are *N*=1000 and *ω*^*2*^=1.

For predominantly selfing populations, the initial dynamics strongly depend on “how much” diversity the population has stored at M-S-D equilibrium (represented by *COV*_LD_). In all cases, additive genetic variance increases during the adaptation process (Figure 5A). When genetic associations are rare (Figure 5, *U* = 0.005), adaptation is driven solely through the increase in frequency of beneficial alleles, leading to an increase of the genic variance and inbred variance (Figures 5B and C). In this case, *COV*_LD_ slightly decreases during the first generations (Figure 5D), probably due to the formation of linkage disequilibrium between alleles rising up in frequency. When among-loci associations are more substantial (*U* > 0.005), the dynamics observed are qualitatively different, and reflect another process of adaptation. The increase in additive variance is due to the “release” of genetic diversity (*COV*_LD_ increases, figure 5D) over the first five generations, whereby the genic and inbred variances remain constant or decrease (Figure 5B and C). The variation of *V*_inbred_ remains equal to *F*.*V*_genic_, indicating that, as observed for outcrossing populations, the inbreeding coefficient does not vary during adaptation.

**Figure 5.**
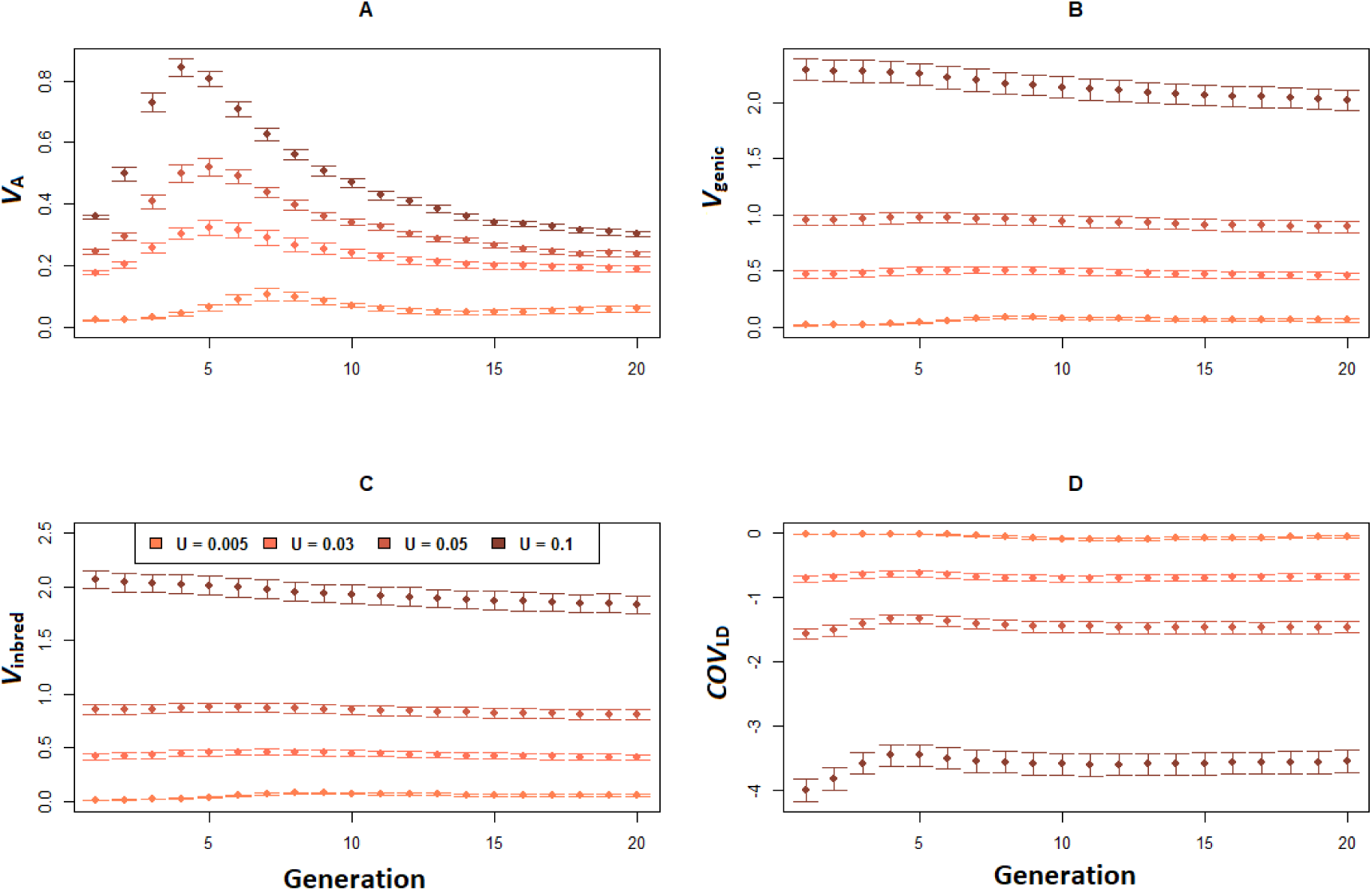
Dynamics of the additive genetic variance and its components in selfing populations (*σ*=0.95), mutation rates are set at zero after generation 0. **A**. Observed additive variance for the phenotypic trait. **B**. Genic variance for the phenotypic trait (*V*_genic_). **C**. Genetic variance due to inbreeding (*V*_inbred_). **D**. Genetic covariance due to linkage disequilibrium (*COV*_LD_), for different mutation rates (U=0.005; U=0.03; U=0.05; U=0.1). Error bars stand for 95% confidence interval (n = 100). Other parameter values are *N*=1000 and *ω*^*2*^=1.

The initial response to selection (increase in trait mean over the first five generations) tightly correlates with the initial amount of additive variance (Figure 6). Nevertheless, for a theoretical equal amount of additive variance at equilibrium (over all parameter sets combined), selfing populations have a stronger initial response at the phenotypic level compared to more allogamous ones (Figure 6). This is due the remobilization of the diversity stored by genetic associations during the first few generations, increasing the additive variance and thus the initial response to selection in selfing populations. The larger the population size, the more the pattern observed for selfing populations differs from that observed in outcrossing and mixed mating ones (Figure 6). When selection is moderate to weak, the release of the stored genetic variation is not necessary for selfing populations to adapt, thus this increase in the strength of adaptation for selfers is no longer observed.

**Figure 6.**
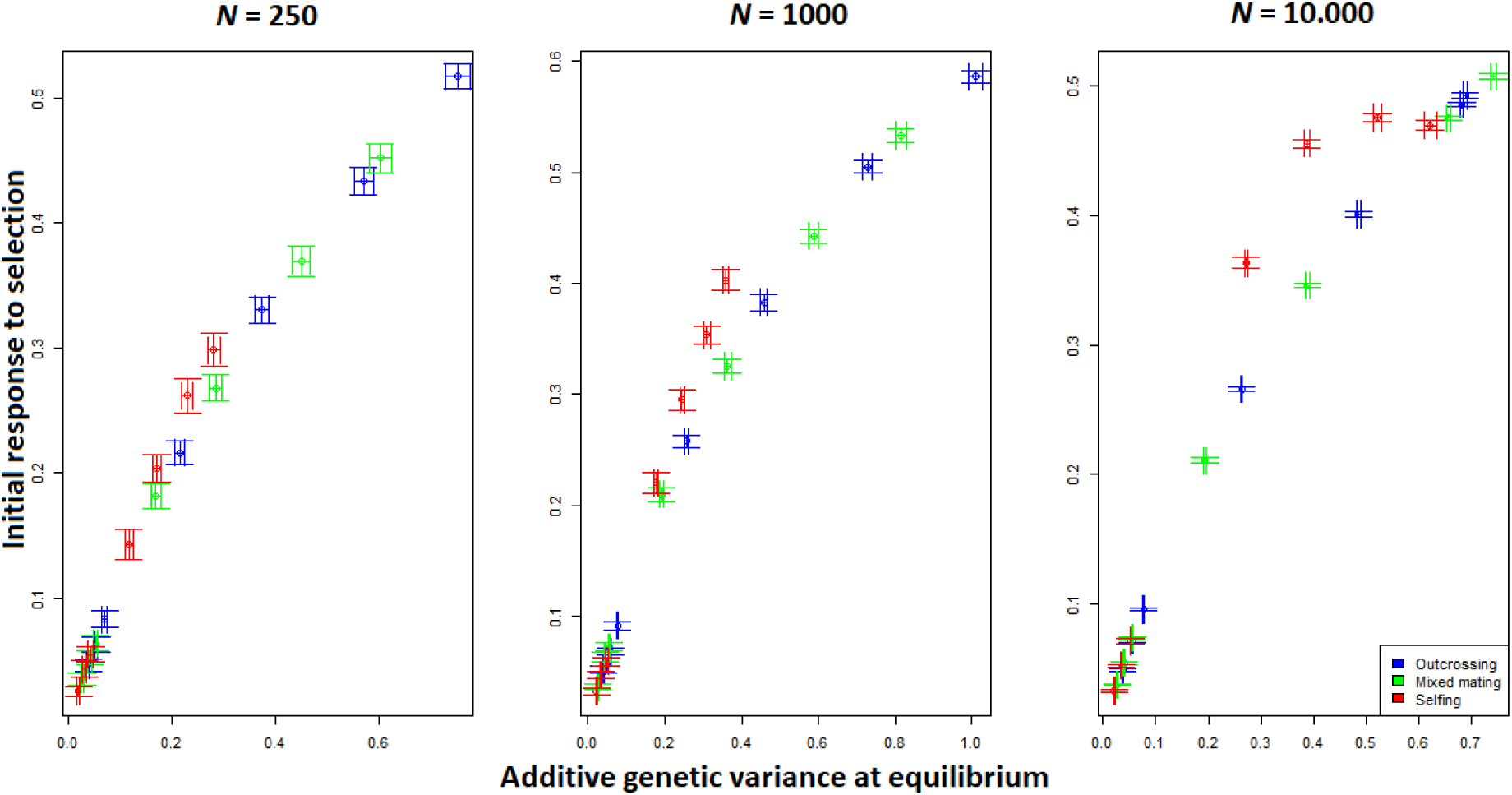
Initial response to selection in function of the amount of additive variance at equilibrium, as a function of the mating system, each point correspond to a haplotypic mutation rate (*U* ranging from 0.005 to 0.1, from the left to the right), and for the three population sizes, when selection is strong (*ω*^*2*^ = 1). Error bars stand for 95% confidence interval (n = 100).

## DISCUSSION

In accordance with Stebbins’ definition of the dead-end hypothesis (Stebbins, 1957), single-locus models predict that adaptation is less likely in selfing populations compared to outcrossing ones, notably due to the reduced standing genetic variation resulting from purging (Glémin & Ronfort, 2013). Considering a polygenic trait, and among loci associations, we find that this is not always the case. Previous works have highlighted that stabilizing selection favours the build-up of associations between loci and, through this, favours the storage of genetic diversity especially in self-fertilising populations (Charlesworth, 1990; Lande & Porcher, 2015; Abu Awad & Roze, 2018). If the selective regime were to change from stabilising to directional selection, due to a strong environmental effect for example, our simulations show that some of this variance can be released through residual allogamy. Residual allogamy is primordial, as selfing populations are organized in multi-locus genotypes. The release of the diversity stored through genetic associations is only possible through rare outcrossing events between two lineages, resulting in fully heterozygote hybrids, which can potentially generate *D*^3^ new genotypes (*D* being the number of differentiated loci among two selfing lineages) after one generation of selfing. Although the strength of the response to selection is positively correlated with the amount of observable standing genetic variation, it is possible to observe similar levels of adaptation in selfing compared to outcrossing or mixed mating populations, for high but realistic mutation rates (Shaw *et al*., 2002). When the strength of selection is moderate to weak (*ω*^2^ = 9 or 99), populations seem to harbour enough genetic variation to quickly and efficiently respond to an environmental change, independently of the self-fertilization rate, and thus without remobilizing any variance accumulated through genetic associations.

### Genetic variance dynamics under directional selection: the role of genetic associations

Few theoretical studies have tried to predict the dynamic of the additive variance for a trait under directional selection (Barton & Turelli, 1987; Keightley & Hill 1989; Bürger, 1993). From these works, under the assumption of random mating and no association between loci, it is predicted that *V*_A_ remains unchanged in the short-term, if population size is small (*N* < 500, Bürger, 1993) and/or if the inverse of the strength of selection, given by the term *ω*^*2*^, is large compared to the amount of available genetic variance (Barton & Turelli, 1987). This prediction is supported by our results for obligate outcrossing populations (Figures S11 to S15), even when some negative linkage disequilibrium contributes significantly to additive variance (Figures 2 & 4). For selfing populations, the hypothesis of linkage equilibrium is no longer valid. As shown in our analysis and known from previous results (Lande & Porcher, 2015; Abu Awad & Roze, 2018), genetic associations between loci are significant in these populations, notably when the mutation rate is high. In this case, the breakup of genetic associations is necessary for adaptation when most of the genetic variance is hidden in among-loci associations, leading to an increase of observed additive variance. The mutation rates for which such associations contribute significantly to the genetic additive variance are in the range of mutation rates observed for phenotypic traits in the literature (Keightley & Bataillon, 2000; Shaw *et al*., 2002; Haag-Liautard *et al*., 2007), which suggests that these associations are probably common in natural populations. The more frequent observations of transgressive segregation (progeny of a cross being outside the parental phenotypic range) in inbred compared to outbred species also give support to this mechanism (Rieseberg *et al*., 1999; Johansen-Morris & Latta, 2006).

### *De novo* mutations *vs*. standing genetic variation: rethinking adaptation in selfing species?

It has been a long accepted paradigm that the advantage procured by selfing is the more rapid fixation of *de novo* beneficial mutations, independently of their dominance coefficient, compared to outcrossing populations, where recessive beneficial mutations can be lost through drift before selection drives them to fixation, a process known as “Haldane’s sieve” (Haldane, 1927). From single locus theory, it is expected that adaptation through new mutations should be more likely in selfing species, and should be more likely than adaptation from standing genetic variation (Glémin & Ronfort, 2013). However, recent works have suggested that the reduced effective recombination rate in selfing populations adds a disadvantage even when it comes to the fixation of new mutations. Unlike what is expected in outcrossing populations, the fixation of beneficial mutations in selfing populations can be hindered if they appear during a selective sweep triggered by a beneficial allele at another locus (Hartfield & Glémin, 2016). This observation, as well as the results presented here, shows that taking interactions between loci into account can strongly modify the expectations from single-locus models. The mutation rate seems to be the major parameter to consider when studying the dynamics of adaptation from a polygenic point of view, independently of whether adaptation occurs from *de novo* mutations (Hartfield & Glémin, 2016) or standing genetic variation.

In our simulations, we have considered a quantitative trait with a simple architecture, assuming pure additivity of genetic effects on the phenotype. Epistasis, and notably its directionality, is known to play a key role in adaptation (Hansen, 2013). Positive epistasis (several loci reinforcing each other’s effects in the direction of selection), inflates the additive variance and thus the ability of a population to adapt to an environmental change (Carter *et al*., 2005; Monnahan & Kelly, 2015). On the contrary, negative epistasis (loci suppressing effects at other loci), reduces the additive variance of the character, thus limiting adaptive potential (Carter *et al*., 2005). Few empirical estimations of the directionality of epistasis are available in the literature (Le Rouzic 2014; Monnahan and Kelly 2015; Oakley *et al*. 2015, all detecting positive epistatic interactions), despite numerous methods and the diversity of data used to infer it (Le Rouzic, 2014). How the directionality of epistatic interactions varies in relation to the mating system remains unknown in natural populations. Knowledge on this relationship may bring us closer to the understanding of the differences in patterns of adaptation observed between selfing and outcrossing populations.

### New insights into the role of standing genetic variation in the adaptation dynamics of selfing populations

The overwhelming success of selfing species in the domestication process and as invasive species has been attributed to mechanisms other than their adaptive ability from standing genetic variation. For instance, the invasive success of selfing populations is attributed to their reproductive assurance, since a single individual is able to colonize a new environment (Rambuda & Johnson, 2004; van Kleunen *et al*., 2008), and to reduced gene flow, which is expected to limit maladapted gene exchanges between populations (Levin, 2010). Regarding domestication as an adaptation process, it has been suggested that domestication in selfing populations most probably relied on new mutations, due to the initially low genetic variance that would have been further reduced due to the bottleneck effect of domestication (Glémin & Bataillon, 2009). This idea is reinforced by the fact that selfing species are expected to quickly fix a rare beneficial mutation, independently of its dominance level (Ross-Ibarra, 2005). In their review on mating system variation in domesticated plant species, Glémin and Bataillon (2009) have suggested that the high frequency of self-fertilizing crop species could be related to an increase in the amount of additive variance during the domestication process of selection. This idea has, however, never been tested theoretically or empirically. Here we show that this increase in additive variance could indeed be an advantage when selfing species are faced with strong directional selection and selection occurs through standing genetic variation. However, our results hold true only if the bottleneck related to domestication (or other invasion processes) is not too strong and if mutation rates are high enough to maintain enough diversity that can promote adaptation to new conditions.

## CONCLUSION AND PERSPECTIVES

In this work, we show that under stabilizing selection and if mutation rates are high enough (but realistic), selfing populations are able to accumulate genetic variation through negative linkage disequilibrium. This structure of genetic variation contributes to acting as a store of genetic variability, thanks to which, adaptation under high self-fertilisation rates is no longer constrained by the reduced observable additive genetic variance due to purging. For low mutation rates, the storage of diversity is impeded by stabilizing selection in selfing populations, leading to the more classical conclusion that adaptation in selfing species is limited by the available amount of genetic diversity. Our analysis thus shows that measuring the amount of additive variance available for quantitative traits is not always sufficient to make predictions about the adaptive potential of a population. Complementary analyses should be carried out when quantifying the short-term adaptability of a population. Such analyses could include looking for transgressive segregations or carrying out experimental evolutionary experiments in which directional selection is induced, and follow the dynamic of additive genetic variance. Following the components of additive variance is complicated, as it necessitates thorough knowledge of the genetic architecture of the quantitative trait under study (number of loci underlying phenotypic variation, their frequency, the linkage disequilibrium…), which is complicated, even for model species. More empirical evidence is required to determine how frequently diversity is stored through genetic associations in natural populations of selfing species, and whether this property is sufficient to allow selfing species to adapt to a changing environment.

## Supporting information

Sup Mat

## Acknowledgment

We would like to thank D. Roze for helpful discussions about the analyses and interpretations of the results, and E. Noël, P. David, O. Ronce, S. Glémin and C. Devaux for insightful discussions on the results. We also thanks editors and reviewers for improving the manuscript. This work has been conducted with the help of the data and calculation center South Green of the CIRAD-UMR AGAP. This work was supported by project SEAD (ANR-13-ADAP-0011), the TUM University Foundation Fellowship and the Alexander von-Humboldt Foundation.

## Appendix A Impact of the mating system on genetic variance at mutation-selection-drift equilibrium: comparison of simulation results with analytic predictions

Several theoretical works have been developed to explicitly quantify the effect of selfing on the amount of additive genetic variance maintained in a population (Lande, 1977; Charlesworth & Charlesworth, 1995; Lande & Porcher, 2015; Abu Awad & Roze, 2018). However, these approaches often do not take the effects of genetic drift due to small population sizes into account. The Stochastic House of Cards approximation (SHC) provided in Bürger *et al*., (1989), an extension to Turelli’s (1984) House of Cards, explicitly considers the effective population size *N*_E_, which can be chosen so as to reflect the population’s reproductive strategy (see Bürger *et al*., 1989, for the case of dieocious reproduction). Considering these different constructions of models, we briefly compare the analytical expectations of some of the above-mentioned works to our simulation results at mutation-selection-drift equilibrium.

In the SHC framework (Bürger *et al*. 1989), the amount of additive variance at equilibrium is given by:

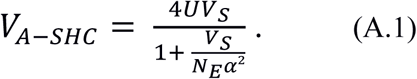

Where *U, V*_S_ and *α* are the haploid mutation rate, the strength of selection and the variance in mutational effects, respectively. Following Caballero & Hill (1992), and ignoring background selection and linkage disequilibrium, the effective population size of an inbreeding population is *N*_E =_ *N*/(1+*F*). *F* is the inbreeding coefficient, which, for a given selfing rate σ, is equal to σ/(2-σ). As mentioned above, this approximation takes into account the population and the strength of selection. However the inbreeding coefficient *F* does not account for consequences of background selection on *N*_E_, which are not known in this specific class of population genetics models (i.e. quantitative traits and stabilising selection) and could thus limit the accuracy of the SHC approximation.

Abu Awad & Roze (2018) provided approximations for the expected level of *V*_A_ under the continuum of alleles model, neglecting genetic drift and associations between loci:

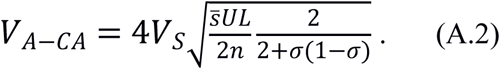

*n* is the number of quantitative traits under consideration (*n* = 1 in our case). In order to account for cases in which linkage is complete (i.e. for very high self-fertilisation rates), setting the number of loci *L* to 1 gives

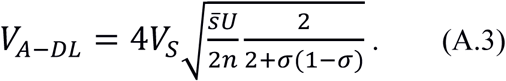

This whole genome is thus modelled as a single locus, with a large number of possible alleles, whose effects follow a Gaussian distribution (Lande, 1977).

To determine the accuracy of these equations under our model conditions, we ran one hundred repetitions per parameter set for three selfing rate values (*σ* = 0, 0.5 and 1), as well as for the seven mutation rates tested in the main text. The mean additive variance from the simulation runs are then compared to the expectations from equations A.1, A.2 and A.3, by using the sum of absolute values of the differences between observations and expectations over the seven mutation rates. Results are presented in Table A1.

**Table A1.** Comparison of analytical predictions for the amount of *V*_A_ under different quantitative genetics models and results obtained in our simulations at MSD equilibrium, for different sets of parameters. The best model for the prediction of *V*_A_ (i.e. closest to our simulation results) is highlighted in green for each parameter set.

| Parameters |  | N = 250 |  |  | N = 1000 |  |  | N= 10 000 |  |  |
| --- | --- | --- | --- | --- | --- | --- | --- | --- | --- | --- |
| | | $V_S$ | | | $V_S$ | | | $V_S$ | | |
|  |  | 1 | 9 | 99 | 1 | 9 | 99 | 1 | 9 | 99 |
| $V_{A-SHC}$ | $\sigma=0$ | 0.16 | 0.89 | 1.56 | 0.49 | 1.92 | 14.58 | 0.31 | 4.38 | 67.81 |
| | $\sigma=0.5$ | 0.11 | 0.28 | 2.81 | 0.18 | 2.05 | 4.71 | 0.22 | 4.65 | 57.87 |
| | $\sigma=0.95$ | 0.80 | 1.76 | 1.13 | 0.86 | 4.10 | 8.10 | 0.36 | 4.93 | 56.71 |
| $V_{A-CA}$ | $\sigma=0$ | 3.49 | 7.16 | 28.57 | 2.94 | 5.11 | 16.93 | 3.28 | 5.95 | 14.66 |
| | $\sigma=0.5$ | 3.92 | 7.34 | 26.89 | 3.16 | 5.55 | 16.53 | 3.14 | 5.65 | 12.74 |
| | $\sigma=0.95$ | 4.61 | 9.79 | 31.77 | 4.30 | 8.47 | 24.37 | 3.59 | 6.55 | 16.29 |
| $V_{A-DL}$ | $\sigma=0$ | 1.30 | 3.52 | 5.57 | 1.85 | 5.57 | 16.85 | 1.51 | 4.74 | 19.12 |
| | $\sigma=0.5$ | 0.95 | 2.75 | 5.21 | 1.39 | 4.52 | 15.31 | 1.42 | 4.42 | 19.06 |
| | $\sigma=0.95$ | 0.25 | 0.86 | 2.17 | 0.53 | 2.09 | 9.02 | 1.23 | 4.01 | 17.01 |

From Table 1, we find that for outcrossing and mixed-mating populations, the Stochastic House of Cars approximation (equation A.1) is the best predictor of the mean additive genetic variance for a wide range of parameters for outcrossing and mixed mating populations. The continuum of alleles approximation (equation A.2) is accurate only for very large population sizes and weak selection, conditions for which the underlying assumptions made when deriving this approximation are met (low allelic frequencies, weak genetic drift and few among-loci associations). For predominantly selfing populations, the approximation considering full linkage (equation A.3) is accurate in conditions where strong genetic associations are maintained throughout the genome *i*.*e*. selection is strong or moderate (table A1).

