## Supplementary material for "Hidden genetic variance contributes to increase the short-term adaptive potential of selfing populations": Sup Mat

**Supplementary materials :**

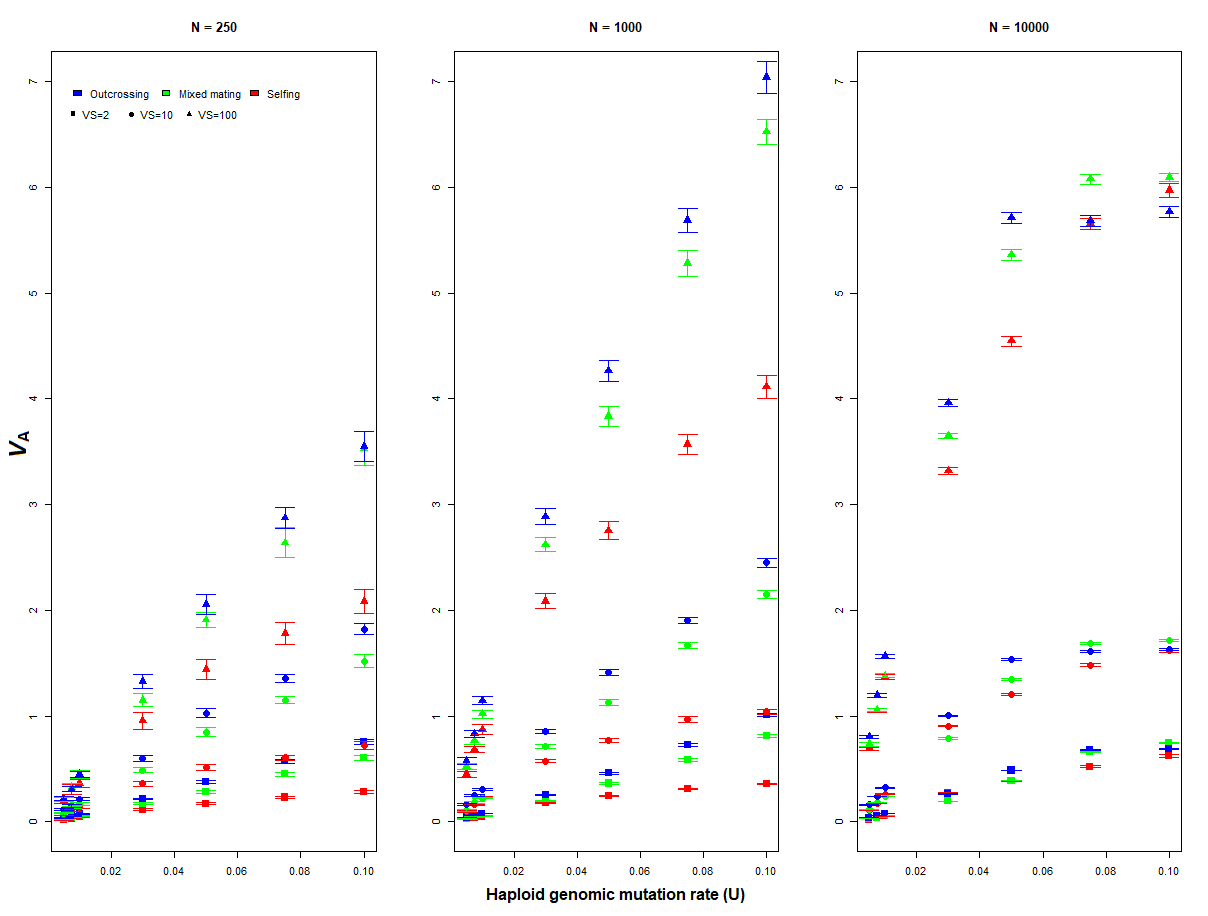

**Figure S1.** Additive genetic variance maintained at mutation-selection-drift equilibrium, in function of the mating system, the population size, and the strength of stabilizing selection. Error bars stand for 95% confidence interval (n = 100).

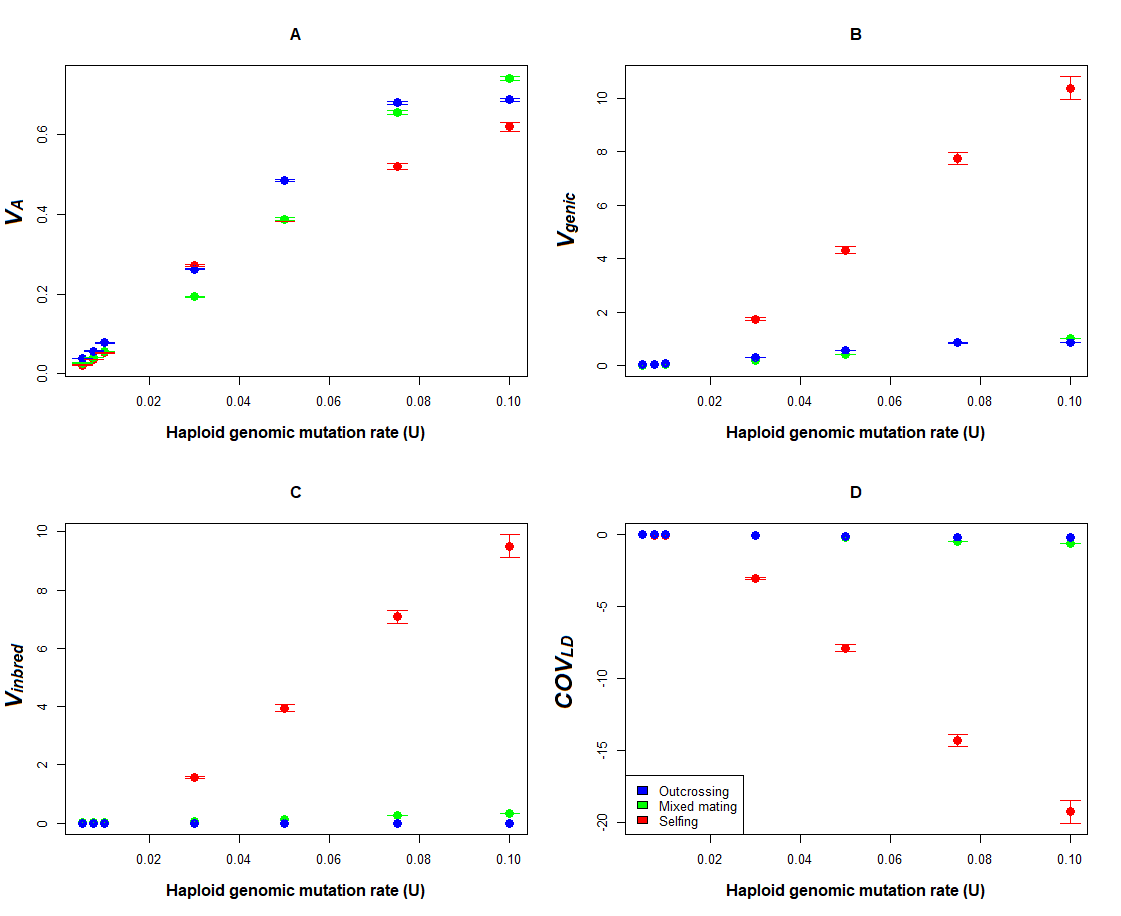

**Figure S2.** Additive genetic variance and its components as a function of the genomic mutation rate and the mating system, for *N*=10.000 and *ω²*=1. **A.** Observed additive variance for the phenotypic trait. **B.** Genic variance for the phenotypic trait (*V*_genic_). **C.** Genetic variance due to inbreeding (*V*_inbred_). **D.** Genetic covariance due to linkage disequilibrium (*COV*_LD_). Error bars stand for 95% confidence interval (n = 100).

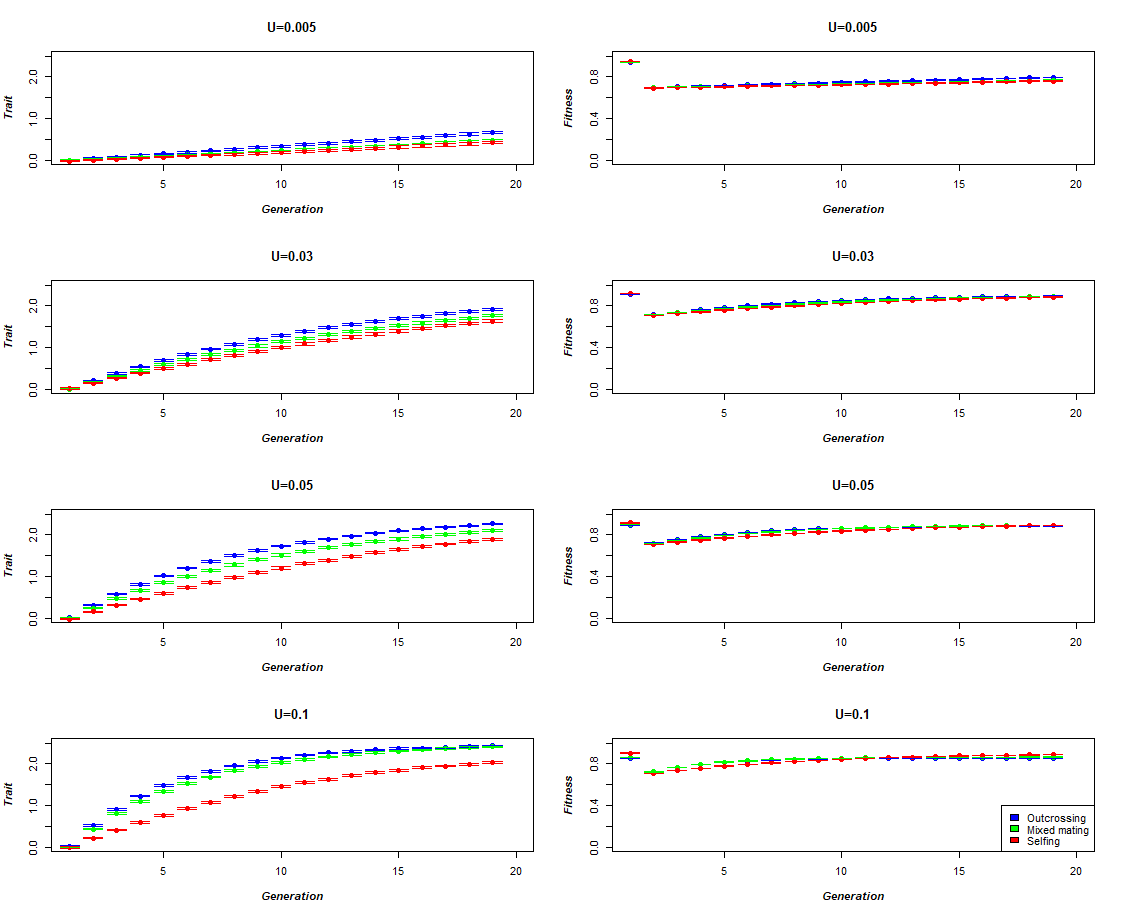

**Figure S3.** Dynamics of the trait and fitness, as a function of the trait haploid mutation rate and the mating system, for *N*=1000 and *ω²*=9. Error bars stand for 95% confidence interval (n=100).

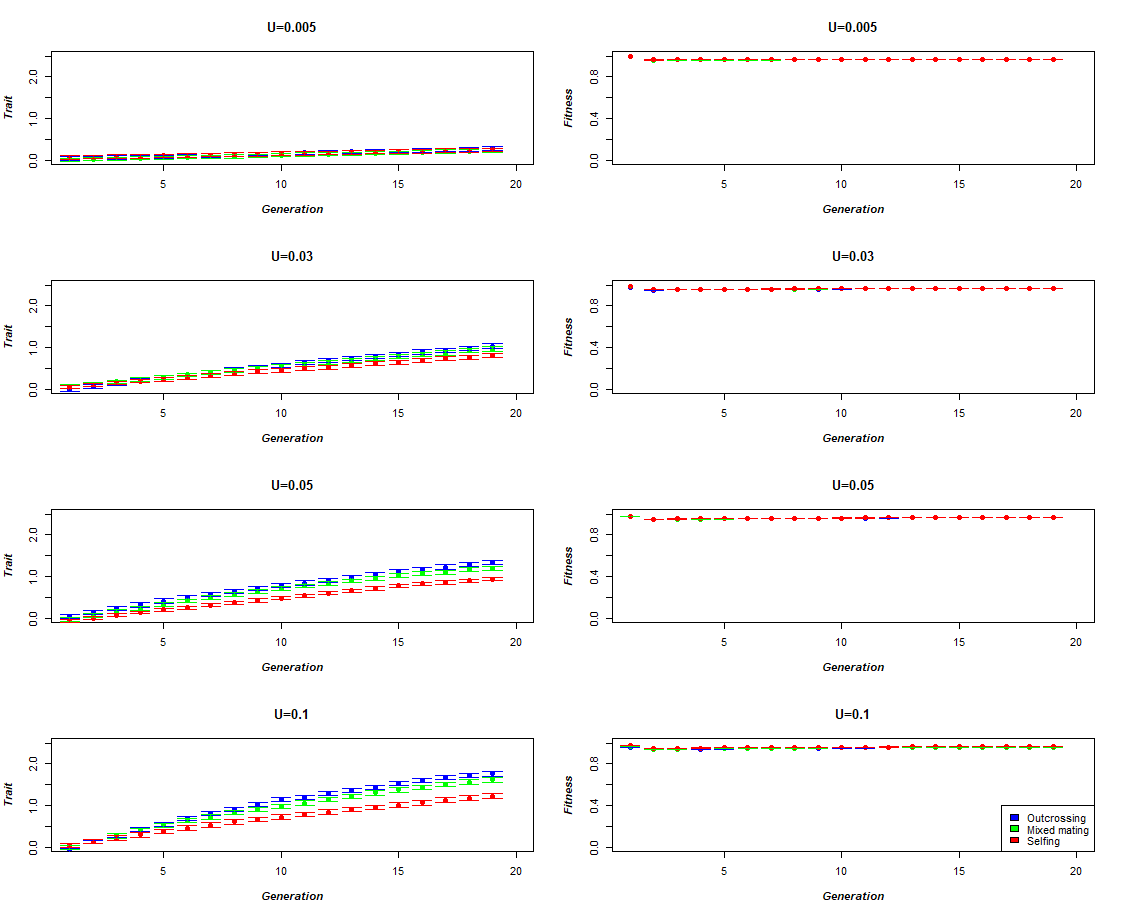

**Figure S4.** Dynamics of the trait and fitness, as a function of the trait haploid mutation rate and the mating system, for *N*=1000 and *ω²*=99. Error bars stand for 95% confidence interval (n=100).

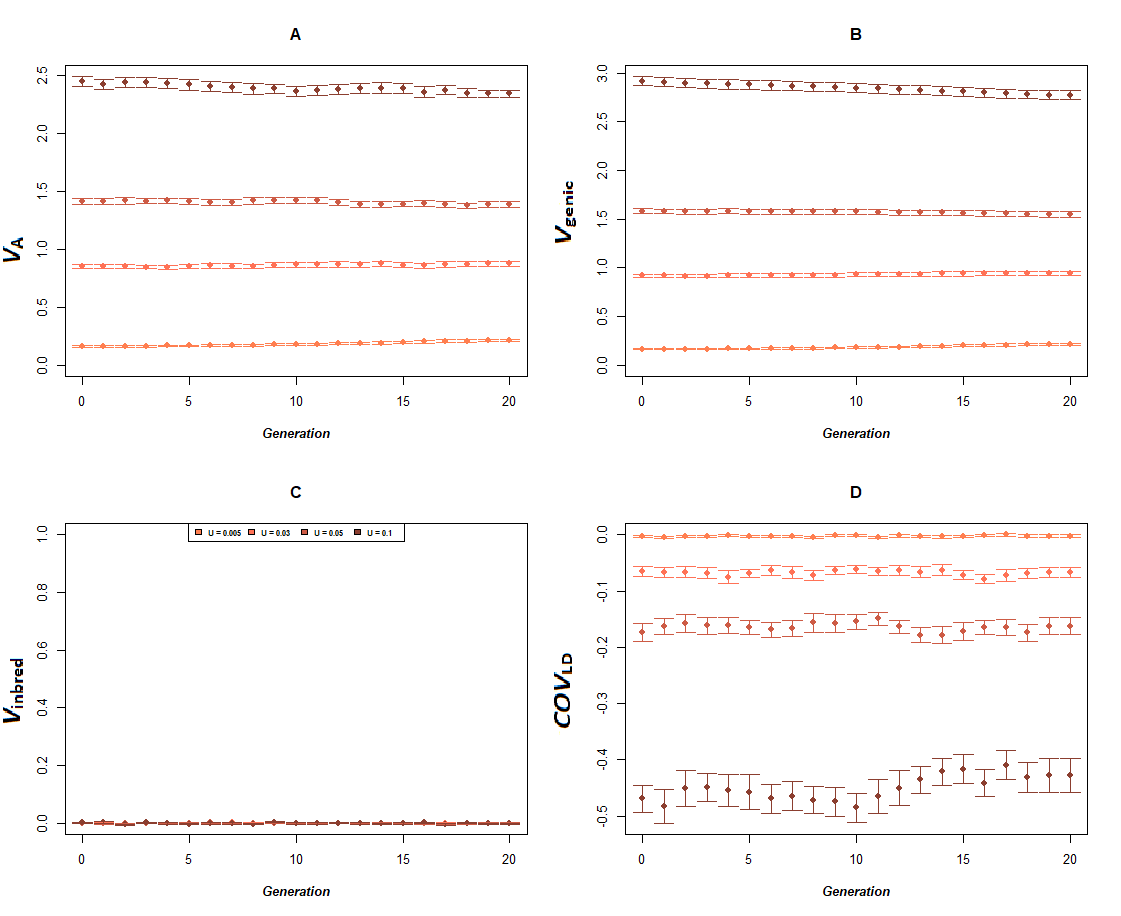

**Figure S5.** Dynamics of additive genetic variance and its components in function of the haplotypic trait mutation rate, for outcrossing populations of *N*=1000 and *ω²*=9. **A.** Observed additive variance for the phenotypic trait. **B.** Genic variance for the phenotypic trait (*V*_genic_). **C.** Genetic variance due to inbreeding (*V*_inbred_). **D.** Genetic covariance due to linkage disequilibrium (*COV*_LD_). Error bars stand for 95% confidence interval (n = 100).

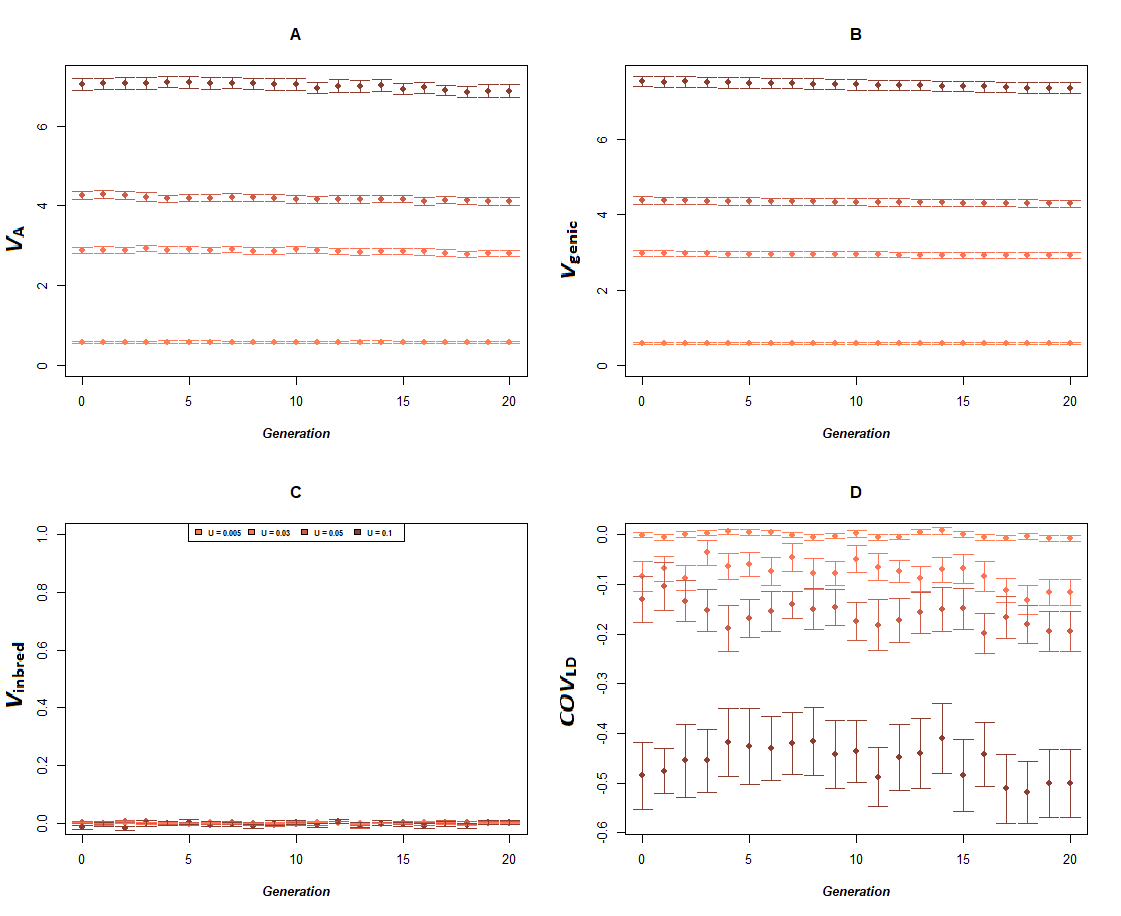

**Figure S6.** Dynamics of additive genetic variance and its components in function of the haplotypic trait mutation rate, for outcrossing populations of *N*=1000 and *ω²*=99. **A.** Observed additive variance for the phenotypic trait. **B.** Genic variance for the phenotypic trait (*V*_genic_). **C.** Genetic variance due to inbreeding (*V*_inbred_). **D.** Genetic covariance due to linkage disequilibrium (*COV*_LD_). Error bars stand for 95% confidence interval (n = 100).

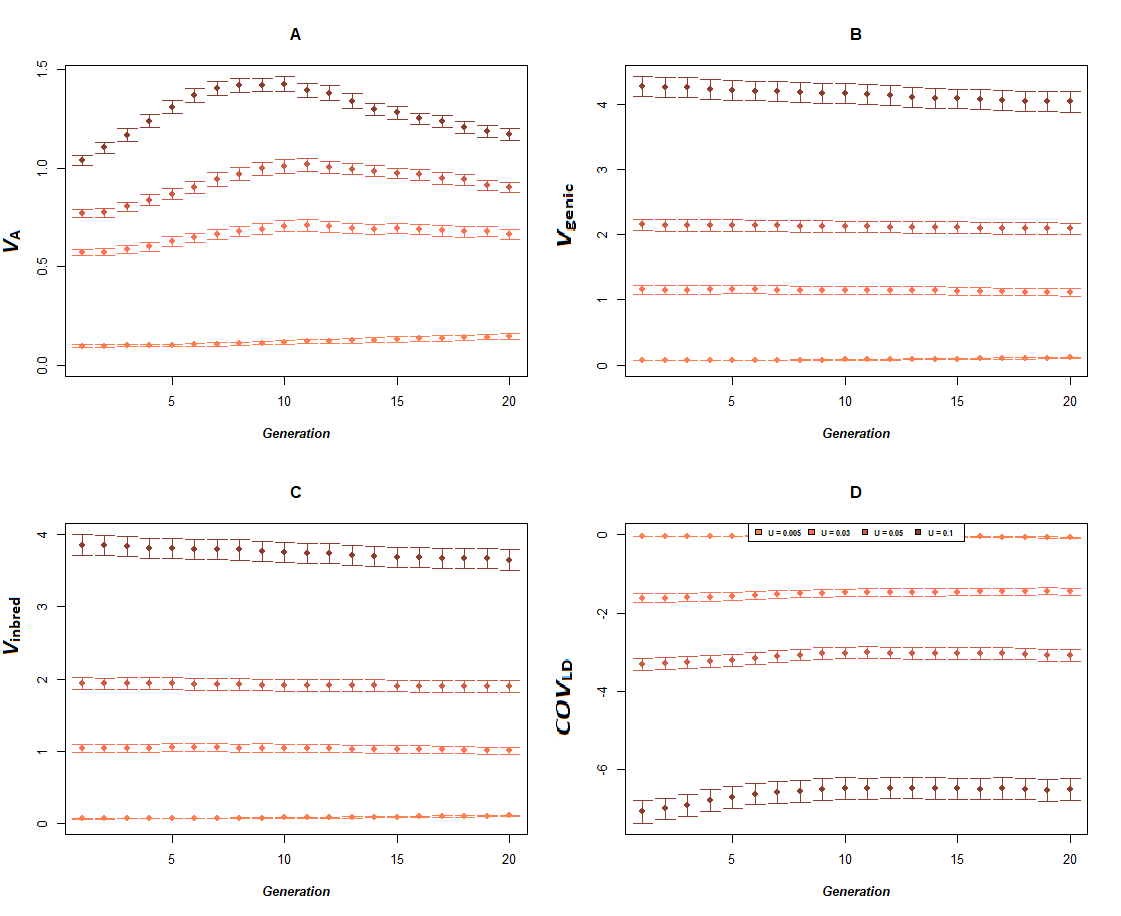

**Figure S7.** Dynamics of additive genetic variance and its components in function of the haplotypic trait mutation rate, for selfing populations of *N*=1000 and *ω²*=9. **A.** Observed additive variance for the phenotypic trait. **B.** Genic variance for the phenotypic trait (*V*_genic_). **C.** Genetic variance due to inbreeding (*V*_inbred_). **D.** Genetic covariance due to linkage disequilibrium (*COV*_LD_). Error bars stand for 95% confidence interval (n = 100).

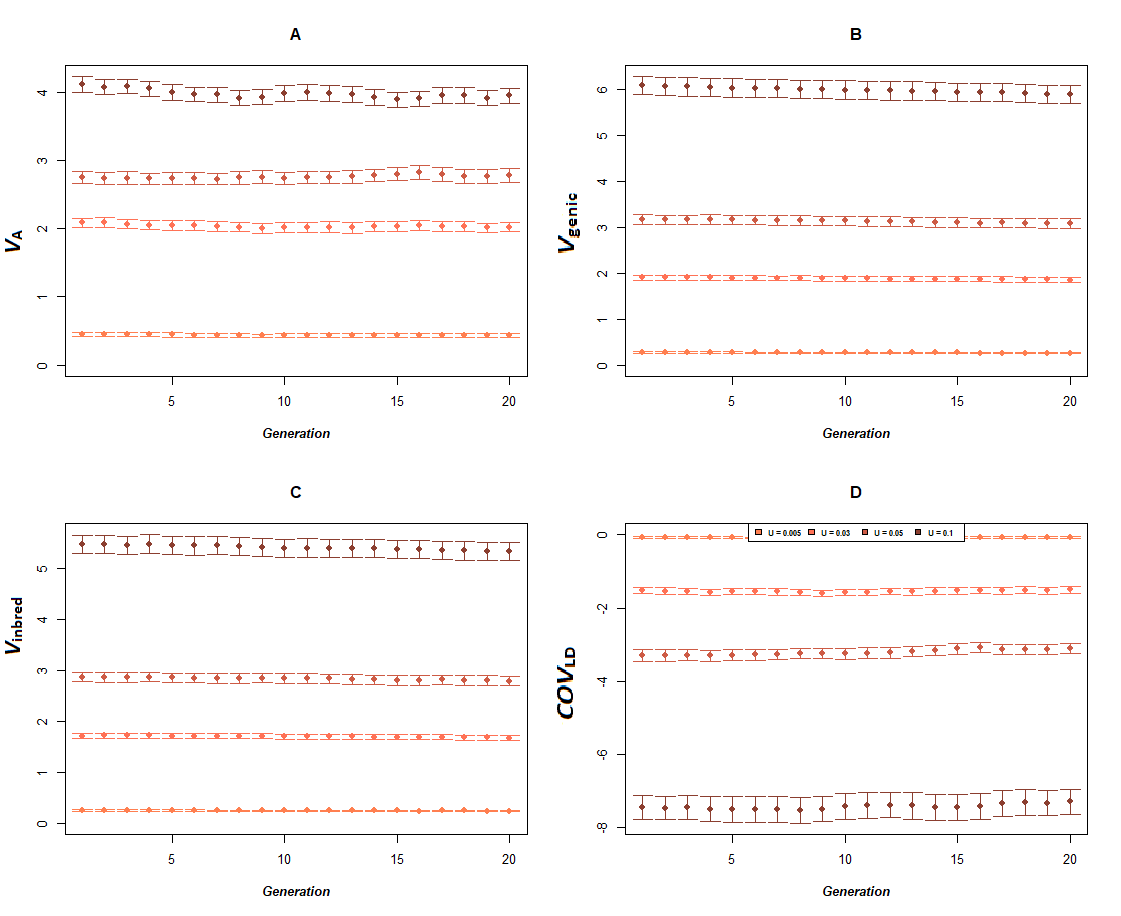

**Figure S8.** Dynamics of additive genetic variance and its components in function of the haplotypic trait mutation rate, for selfing populations of *N*=1000 and *ω²*=99. **A.** Observed additive variance for the phenotypic trait. **B.** Genic variance for the phenotypic trait (*V*_genic_). **C.** Genetic variance due to inbreeding (*V*_inbred_). **D.** Genetic covariance due to linkage disequilibrium (*COV*_LD_). Error bars stand for 95% confidence interval (n = 100).

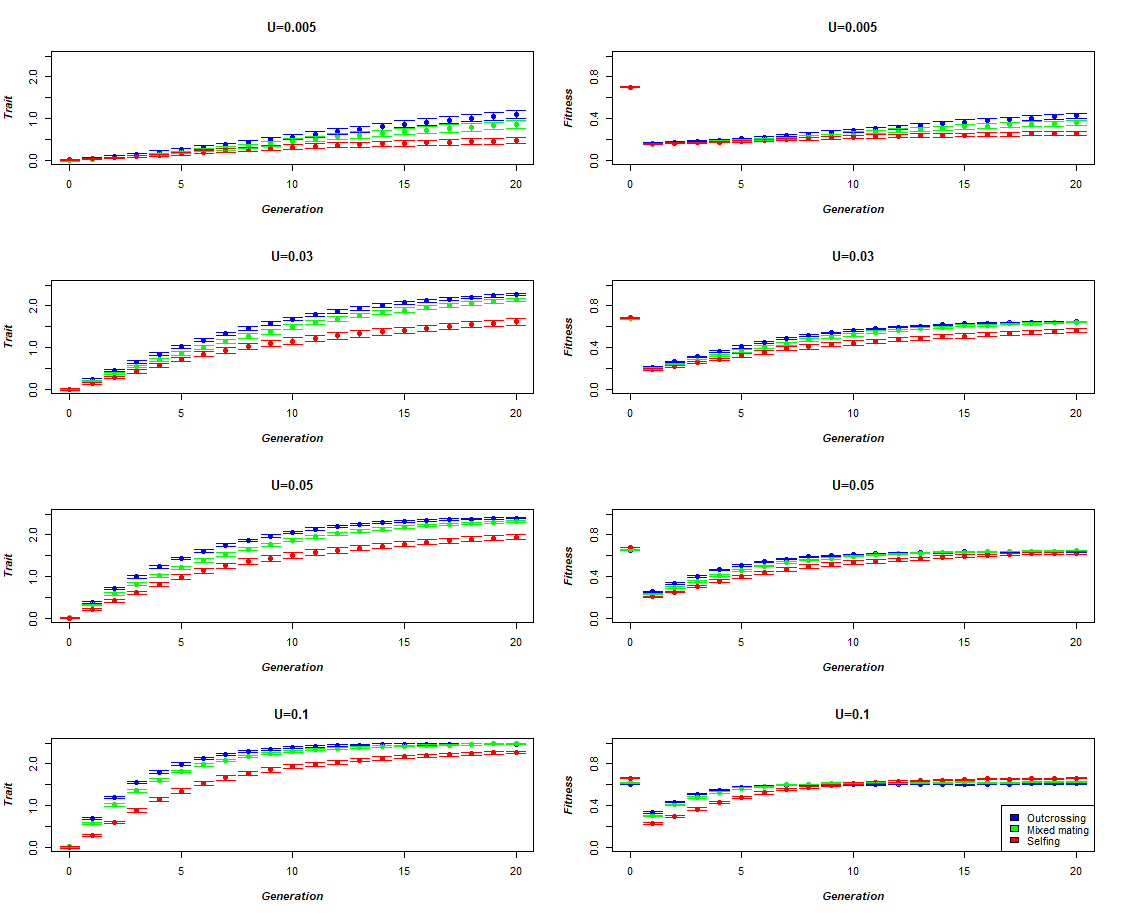

**Figure S9.** Dynamics of the trait and fitness, as a function of the trait haploid mutation rate and the mating system, for *N*=250 and *ω²*=1. Error bars stand for 95% confidence interval (n=100).

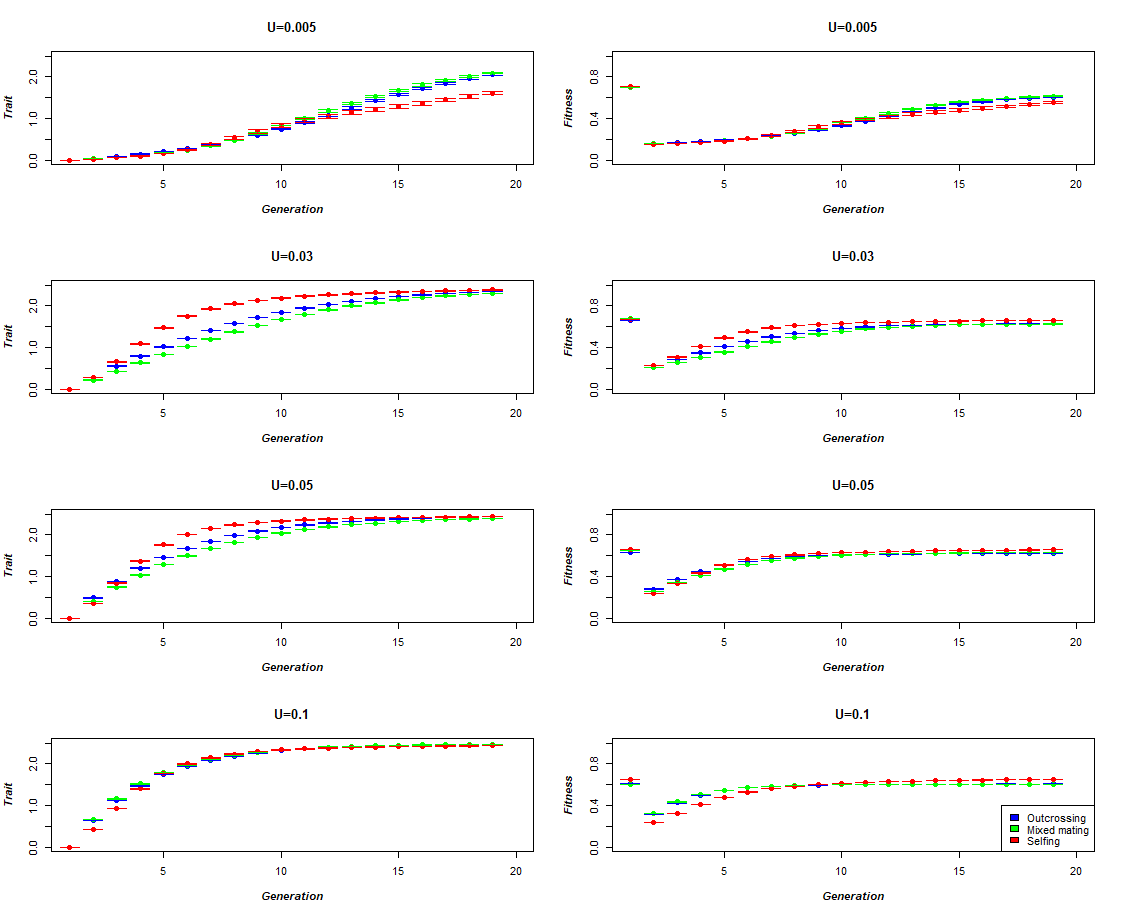

**Figure S10.** Dynamics of the trait and fitness, as a function of the trait haploid mutation rate and the mating system, for *N*=10.000 and *ω²*=1. Error bars stand for 95% confidence interval (n=100).

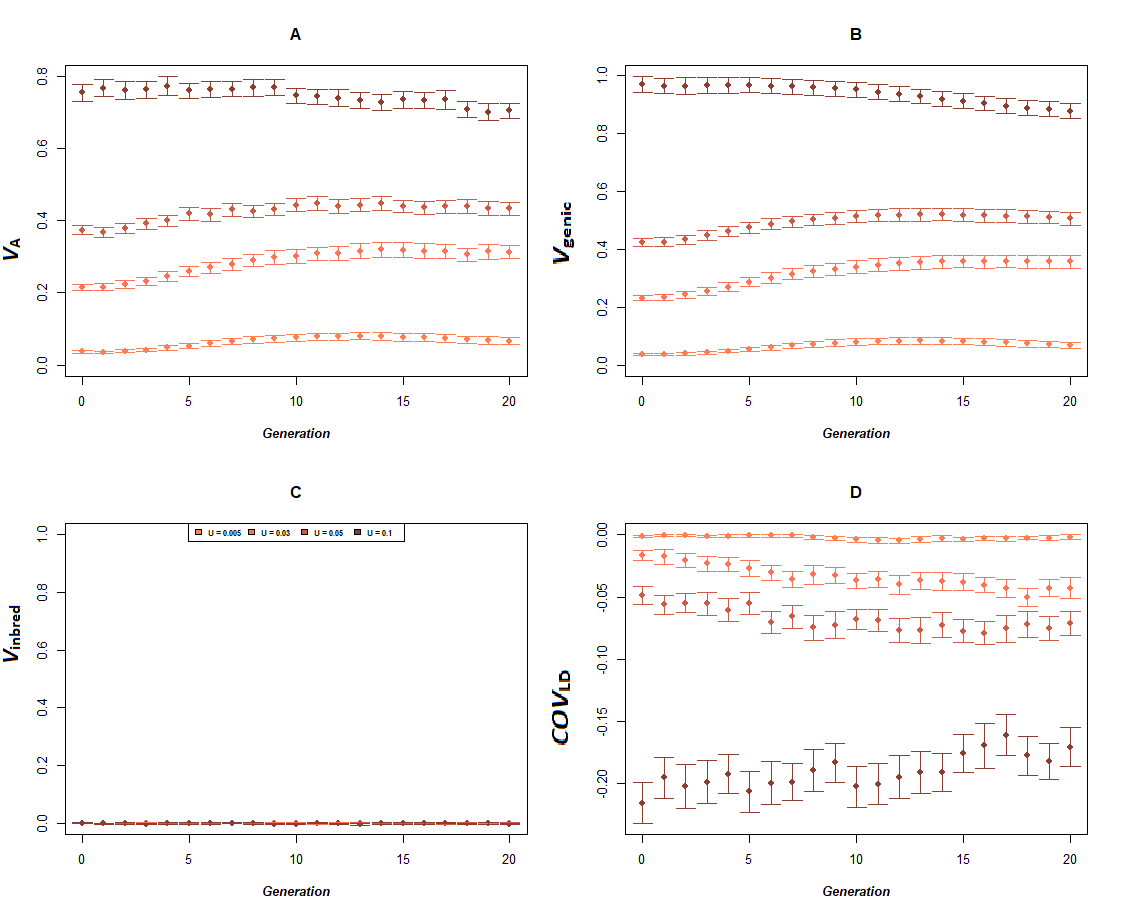

**Figure S11.** Dynamics of additive genetic variance and its components in function of the haplotypic trait mutation rate, for outcrossing populations of *N*=250 and *ω²*=1. **A.** Observed additive variance for the phenotypic trait. **B.** Genic variance for the phenotypic trait (*V*_genic_). **C.** Genetic variance due to inbreeding (*V*_inbred_). **D.** Genetic covariance due to linkage disequilibrium (*COV*_LD_). Error bars stand for 95% confidence interval (n = 100).

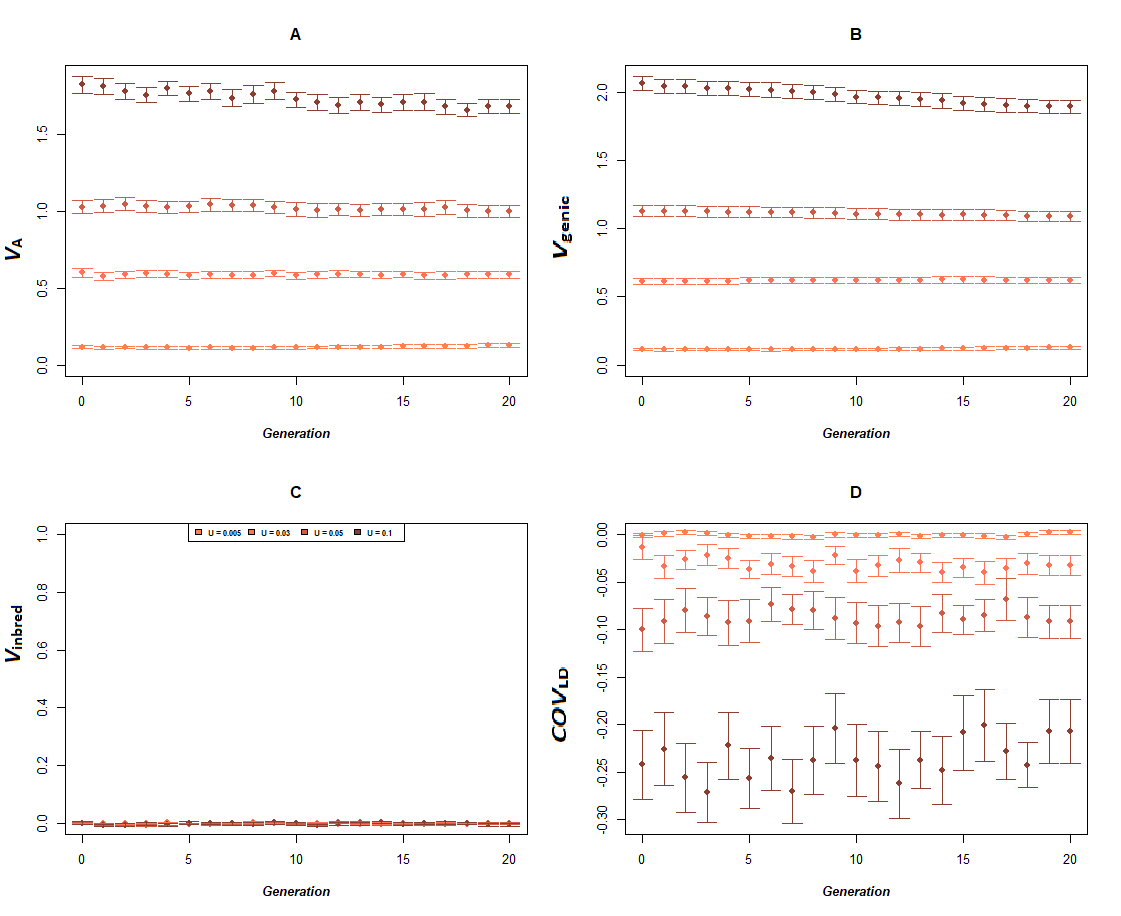

**Figure S12.** Dynamics of additive genetic variance and its components in function of the haplotypic trait mutation rate, for outcrossing populations of *N*=250 and *ω²*=9. **A.** Observed additive variance for the phenotypic trait. **B.** Genic variance for the phenotypic trait (*V*_genic_). **C.** Genetic variance due to inbreeding (*V*_inbred_). **D.** Genetic covariance due to linkage disequilibrium (*COV*_LD_). Error bars stand for 95% confidence interval (n = 100).

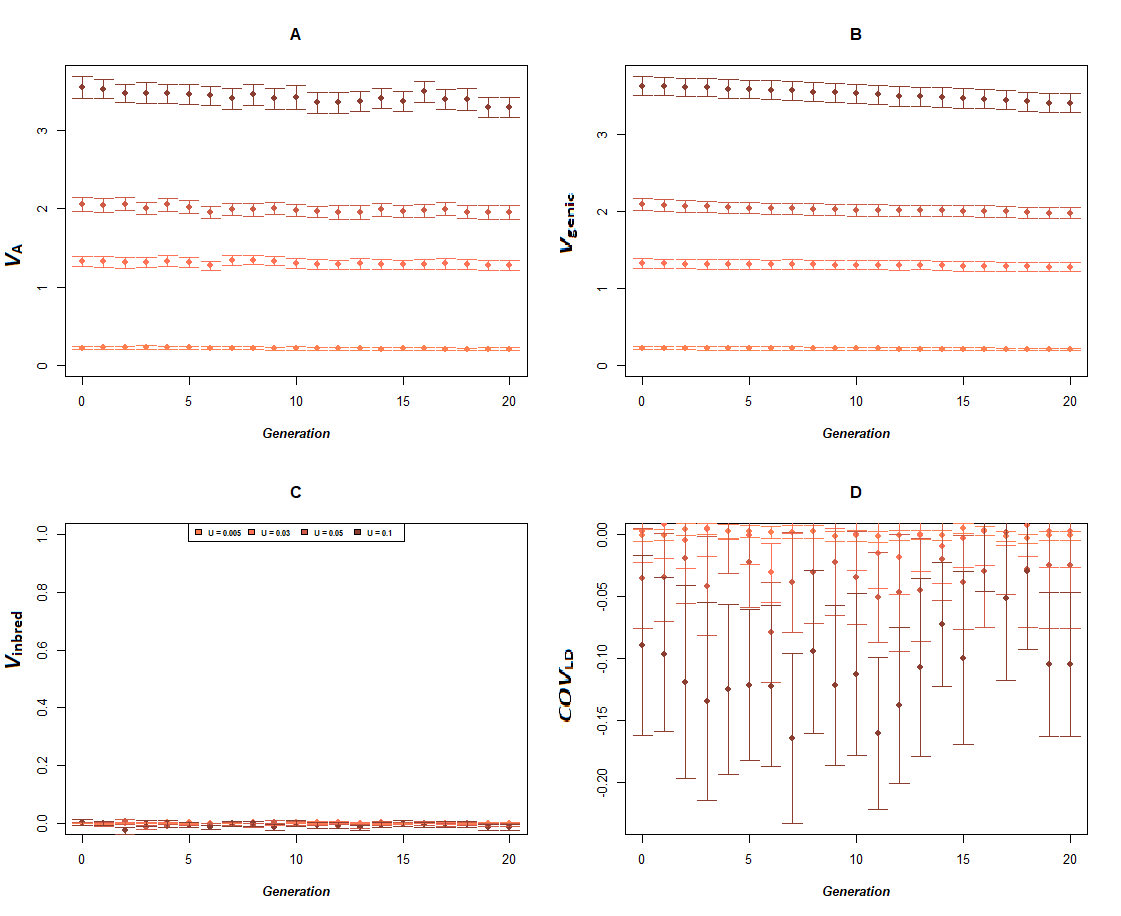

**Figure S13.** Dynamics of additive genetic variance and its components in function of the haplotypic trait mutation rate, for outcrossing populations of *N*=250 and *ω²*=99. **A.** Observed additive variance for the phenotypic trait. **B.** Genic variance for the phenotypic trait (*V*_genic_). **C.** Genetic variance due to inbreeding (*V*_inbred_). **D.** Genetic covariance due to linkage disequilibrium (*COV*_LD_). Error bars stand for 95% confidence interval (n = 100).

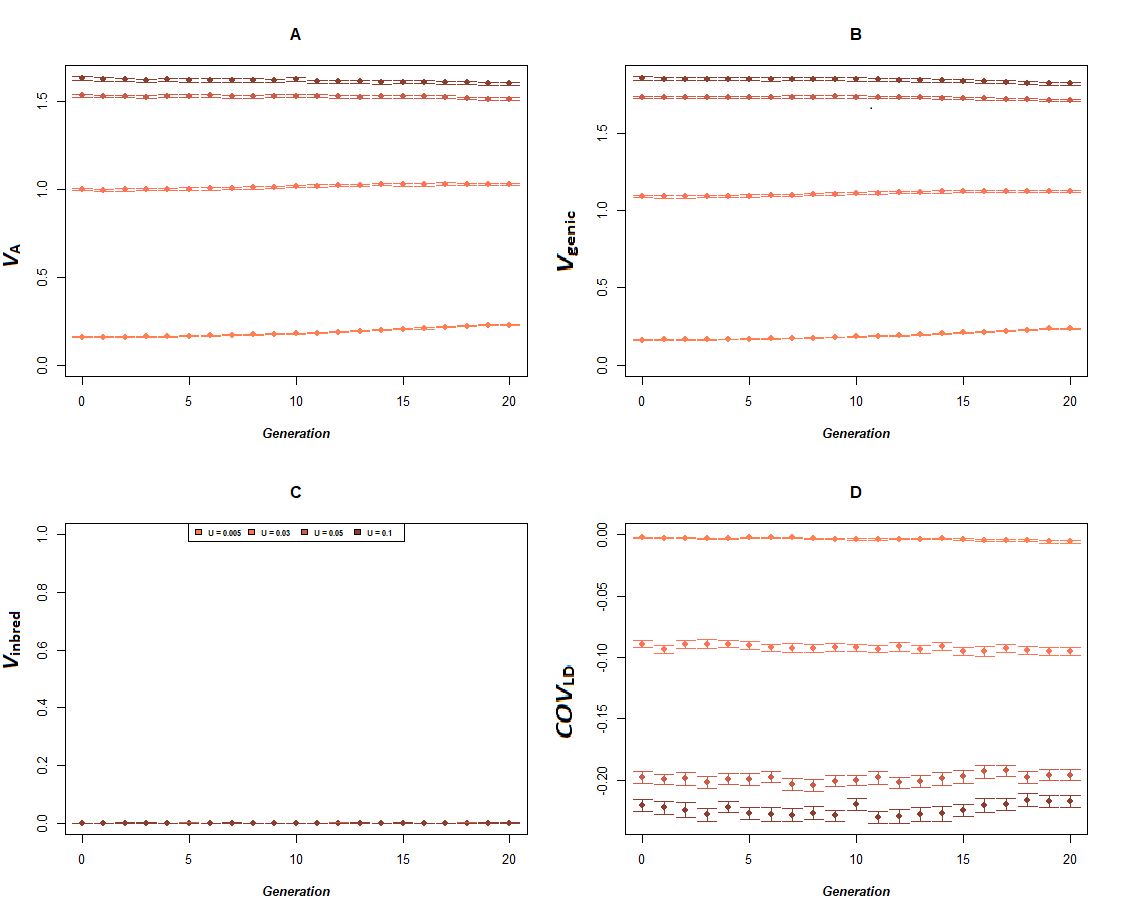

**Figure S14.** Dynamics of additive genetic variance and its components in function of the haplotypic trait mutation rate, for outcrossing populations of *N*=10.000 and *ω²*=9. **A.** Observed additive variance for the phenotypic trait. **B.** Genic variance for the phenotypic trait (*V*_genic_). **C.** Genetic variance due to inbreeding (*V*_inbred_). **D.** Genetic covariance due to linkage disequilibrium (*COV*_LD_). Error bars stand for 95% confidence interval (n = 100).

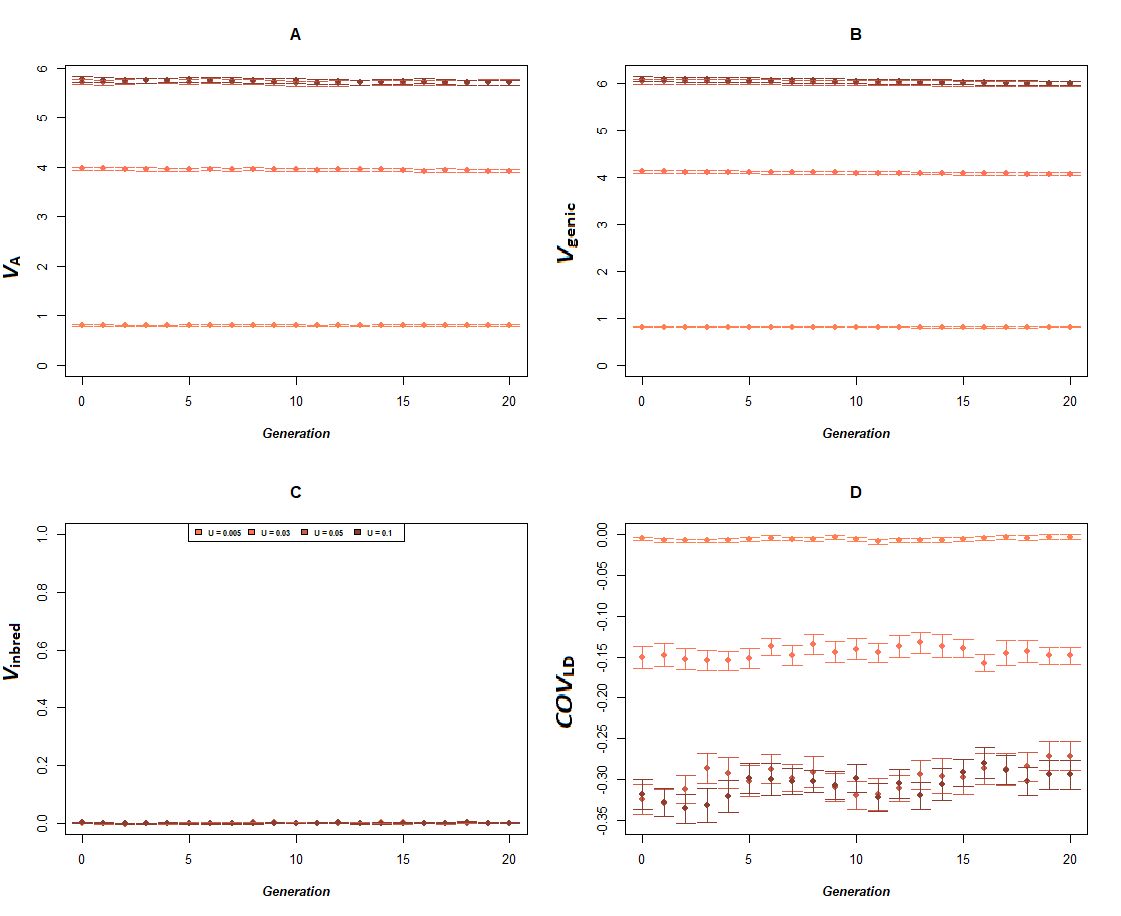

**Figure S15.** Dynamics of additive genetic variance and its components in function of the haplotypic trait mutation rate, for outcrossing populations of *N*=10.000 and *ω²*=99. **A.** Observed additive variance for the phenotypic trait. **B.** Genic variance for the phenotypic trait (*V*_genic_). **C.** Genetic variance due to inbreeding (*V*_inbred_). **D.** Genetic covariance due to linkage disequilibrium (*COV*_LD_). Error bars stand for 95% confidence interval (n = 100).
